# TGF-Β Family Inhibitors Blunt Adipogenesis Via Non-Canonical Regulation Of SMAD Pathways

**DOI:** 10.1101/2020.03.12.988568

**Authors:** Senem Aykul, Jordan Maust, Monique Floer, Erik Martinez-Hackert

## Abstract

Adipose tissues (AT) expand in response to energy surplus through adipocyte hypertrophy and hyperplasia (*i.e.*, adipogenesis). The latter is a process by which multipotent precursors differentiate into mature adipocytes. This process is directed by growth factors and cytokines, including members of the TGF-β family, which regulate intracellular signaling pathways that control adipogenic transcriptional programs. As ectopic adipogenesis has been linked with metabolic syndrome and other pathological conditions, we undertook to establish how TGF-β family growth factors and their inhibitors regulate this process in a 3T3-L1 adipogenesis model. We found that intracellular SMAD1/5/8 signaling pathways are activated while SMAD2/3 pathways are suppressed in differentiating cells. Addition of SMAD1/5/8 pathway activating ligands promoted cell proliferation, while SMAD2/3 pathway activating ligands suppressed adipocyte formation. We identified several ligand traps that blunted 3T3-L1 adipogenesis. Strikingly, anti-adipogenic traps and ligands exploited the same mechanism of regulation involving a negative feedback loop that links SMAD2/3 activation with SMAD1/5/8 hyper-phosphorylation, cytoplasmic retention, and reduced signaling. The identified anti-adipogenic traps could be used to control hyperplastic AT expansion and its associated pathological conditions.

## INTRODUCTION

Adipose tissues (ATs) are essential for regulating energy balance and for maintaining metabolic, endocrine, and immune health (Rutkowski, Stern et al., 2015). In obesity, which is characterized by the massive expansion of AT driven by disproportionately high energy intake relative to energy expenditure, these regulating functions can become severely compromised, triggering pathological conditions such as type 2 diabetes, cardiovascular disease, nonalcoholic steatohepatitis, and cancers (Eckel, Grundy et al., 2005, Kaplan, 1989). Understanding the mechanisms and factors that underlie growth, size, and function of AT is, therefore, critical from a public health point of view and could help in the development of new therapeutic strategies and targets for treating obesity and its associated complications.

AT expansion in response to caloric excess is driven by two distinct developmental processes (Rutkowski et al., 2015). Existing adipocytes can become hypertrophic as they store surplus energy as triglycerides, while AT can undergo hyperplasia (a.k.a. adipogenesis) as new adipocytes develop from resident precursors (Sun, Kusminski et al., 2011, Sun & Scherer, 2010, Wang, Tao et al., 2013). Transforming Growth Factor-β (TGF-β) family ligands, which include TGF-β, Activin, GDF, and BMP family growth factors, have known roles in both adipogenesis and adipocyte hypertrophy (Lee, 2018, Tang & Lane, 2012, Zamani & Brown, 2011). Thus, TGF-β1, GDF-8, GDF-11, and Activins A and B inhibit adipogenesis (Choy, Skillington et al., 2000, Hirai, Yamanaka et al., 2005, Hoggard, Cruickshank et al., 2009, Ignotz & Massague, 1985, Kim, Liang et al., 2001, Lee, Pickering et al., 2019, Luo, Guo et al., 2019, Sparks, Allen et al., 1992), whereas several BMPs have been shown to promote adipogenesis and/or adipocyte hypertrophy (Gustafson, Hammarstedt et al., 2015, Huang, Song et al., 2009, Modica & Wolfrum, 2017, Schreiber, Dorpholz et al., 2017, Tang, Otto et al., 2004, Tseng, Kokkotou et al., 2008). Yet in spite of a large body of work in this area, fundamental questions remain unresolved, including what steps of adipogenesis are regulated by specific TGF-β family pathways, how are intracellular signaling pathways activated to direct adipocyte development, and can extracellular TGF-β family inhibitors help control adipogenesis.

To address these questions, we investigated the roles of 11 TGF-β family ligands and 11 inhibitory ligand traps in precursor commitment, proliferation and adipocyte hypertrophy using the 3T3-L1 adipogenesis model. Confirming earlier results, we found that TGF-β-like ligands, which primarily activate intracellular SMAD2/3 pathways (Derynck & Zhang, 2003, Shi & Massague, 2003), suppressed adipogenesis. By contrast, BMP-like ligands, which primarily activate SMAD1/5/8 pathways (Derynck & Zhang, 2003, Shi & Massague, 2003), promoted adipocyte proliferation. Using a cell-based screening approach, we identified three ligand traps that profoundly suppressed adipocyte differentiation. Strikingly, anti-adipogenic traps and ligands exploited the same mechanism of regulation involving concomitant SMAD2/3 activation and SMAD1/5/8 hyper-phosphorylation, which in turn associated with sharply reduced SMAD1/5/8 nuclear translocation and transcriptional activity. As SMAD1/5/8 signaling was induced and SMAD2/3 signaling was suppressed during adipocyte differentiation, we propose that SMAD1/5/8 signaling primes and drives commitment of 3T3-L1 cells to the adipogenic fate. By contrast, SMAD2/3 signaling arrests pre-adipocytes in an undifferentiated state or reprograms precursors toward a different fate. Consequently, the SMAD1/5/8 signaling inhibitor LDN-193189 suppressed adipogenesis, while simultaneous treatment of differentiating cells with both adipogenesis inhibitor traps or ligands and the SMAD2/3 signaling inhibitor SB-431542 rescued adipogenesis, prevented SMAD1/5/8 hyper-phosphorylation, and restored SMAD1/5/8 signaling, revealing a negative regulatory feedback loop between the two SMAD branches. In conclusion, we have identified three ligand traps that suppressed adipogenesis *in vitro* and we have elucidated their conserved anti-adipogenic mechanism. These ligand traps could help control AT hyperplasia *in vivo*, and thus could be used to limit pathological conditions associated with hyperplastic AT expansion.

## RESULTS

### TGF-β family ligands regulate early adipogenic fates

TGF-β pathways are known to regulate adipogenic fates (Lee, 2018). To identify steps controlled by TGF-β pathways in adipogenesis, we investigated the time-dependent effect of 11 different ligands on 3T3-L1 differentiation following a standard 3T3-L1 differentiation assay (Zebisch, Voigt et al., 2012) (Figure 1A). We found that TGF-β-like ligands (*i.e.* TGF-βs, Activins, GDF-8 and GDF-11, which mainly activate SMAD2/3 pathways via the receptor kinases ALK4, ALK5, or ALK7 (Derynck & Zhang, 2003, Shi & Massague, 2003)), inhibited early steps of 3T3-L1 adipogenesis as indicated by the near complete absence of lipid droplet (LD) formation in samples treated from the beginning of differentiation (Figures 1B,C). This effect was significantly attenuated but still evident when cells were treated at later stages of differentiation. Intriguingly, the few LDs that formed in treated samples accumulated more lipid (Figures 1B,D). In addition, TGF-β-like ligands reduced 3T3-L1 proliferation, as treated samples had fewer nuclei per well than untreated controls (Figures 1E). Together, these results indicate that TGF-β-like ligands inhibit differentiation and proliferation of adipocyte precursors and may promote adipocyte hypertrophy. Although GDF-8 appeared to be an exception, higher concentrations may be required due to its relatively lower potency (Rebbapragada, Benchabane et al., 2003, Walker, Czepnik et al., 2017).

**Figure 1.**
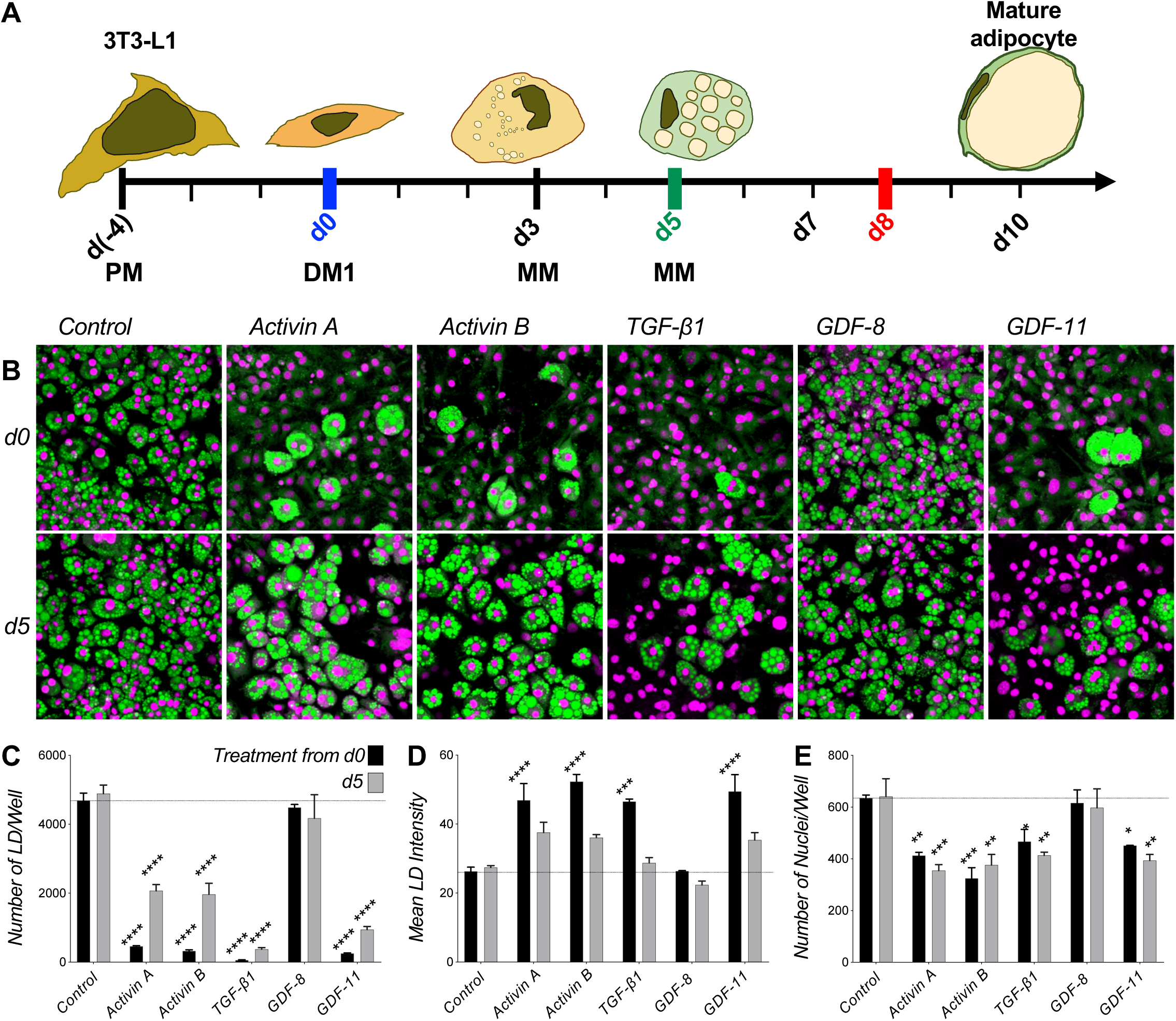
TGF-βs and activins inhibit differentiation of 3T3-L1 cells into adipocytes. **A)** Schematic of 3T3-L1 assay. Cells are grown 4 days in Preadipocyte Medium (PM) (from day −4 to 0), differentiated for 3 days using Differentiation Media, and maintained up to 7 days in Maintenance Medium (MM from day 3 to 10). Cells are treated with ligands or traps either from day 0 (blue) or from day 5 (green) until harvest at day 8 (red). **B)** 3T3-L1 cells were grown with 1 nM ligand as noted (except TGF-β1 at 0.1 nM) or vehicle control (PBS) from day 0 (top panel) or day 5 (bottom panel) of differentiation until day 8. Cells were fixed at day 8 and stained for lipids using Nile red (green), nuclei were counter-stained with DAPI (magenta). Cells treated with this group of ligands mostly showed significantly reduced lipid droplet formation. **C-E)** Quantitative analysis of 3T3-L1 samples treated with ligands from day 0 (black) or day 5 (grey) of differentiation. Images where analyzed with ImageJ and data were evaluated using GraphPad. Statistical significance from two biological replicates was calculated by two-way ANOVA and Dunnett’s multiple comparisons test (* p<0.05; **p<0.01; ***p<0.001, ****p<0.0001). **C)** Total number of lipid droplets (LD) per well. **D)** Mean lipid droplet intensity. **E)** Number of nuclei per well.

In contrast to TGF-β-like ligands, BMP-like ligands (*i.e.* BMPs and most GDFs, which activate SMAD1/5/8 pathways via the receptor kinases ALK1, ALK2, ALK3, or ALK6 (Derynck & Zhang, 2003, Shi & Massague, 2003)) mostly promoted 3T3-L1 proliferation as indicated by the greater number of nuclei per well relative to untreated controls (Figures 2A-D). However, their activities were rather divergent. BMP-2, BMP-4, BMP-6, and BMP-7, which signal via the kinases ALK2, ALK3, or ALK6 (de Caestecker, 2004), promoted proliferation with varying degrees of potency and statistical significance (Figure 2D). By contrast, BMP-9 and BMP-10, which signal predominantly via the receptor kinase ALK1 but also ALK2 (David, Mallet et al., 2007, Herrera, van Dinther et al., 2009, Olsen, Sankar et al., 2018, Olsen, Wader et al., 2014, Scharpfenecker, van Dinther et al., 2007, Townson, Martinez-Hackert et al., 2012), respectively prevented differentiation and promoted proliferation (Figures 2B,D). Notably, BMP-like ligands did not significantly stimulate lipid accumulation as evidenced by the stable number of lipid droplets and lipid droplet size in treated and control samples (Figures 2B,C). Collectively, these findings support a model where BMP-like ligands that signal via the kinases ALK2, ALK3, or ALK6 mainly promote adipocyte proliferation.

**Figure 2.**
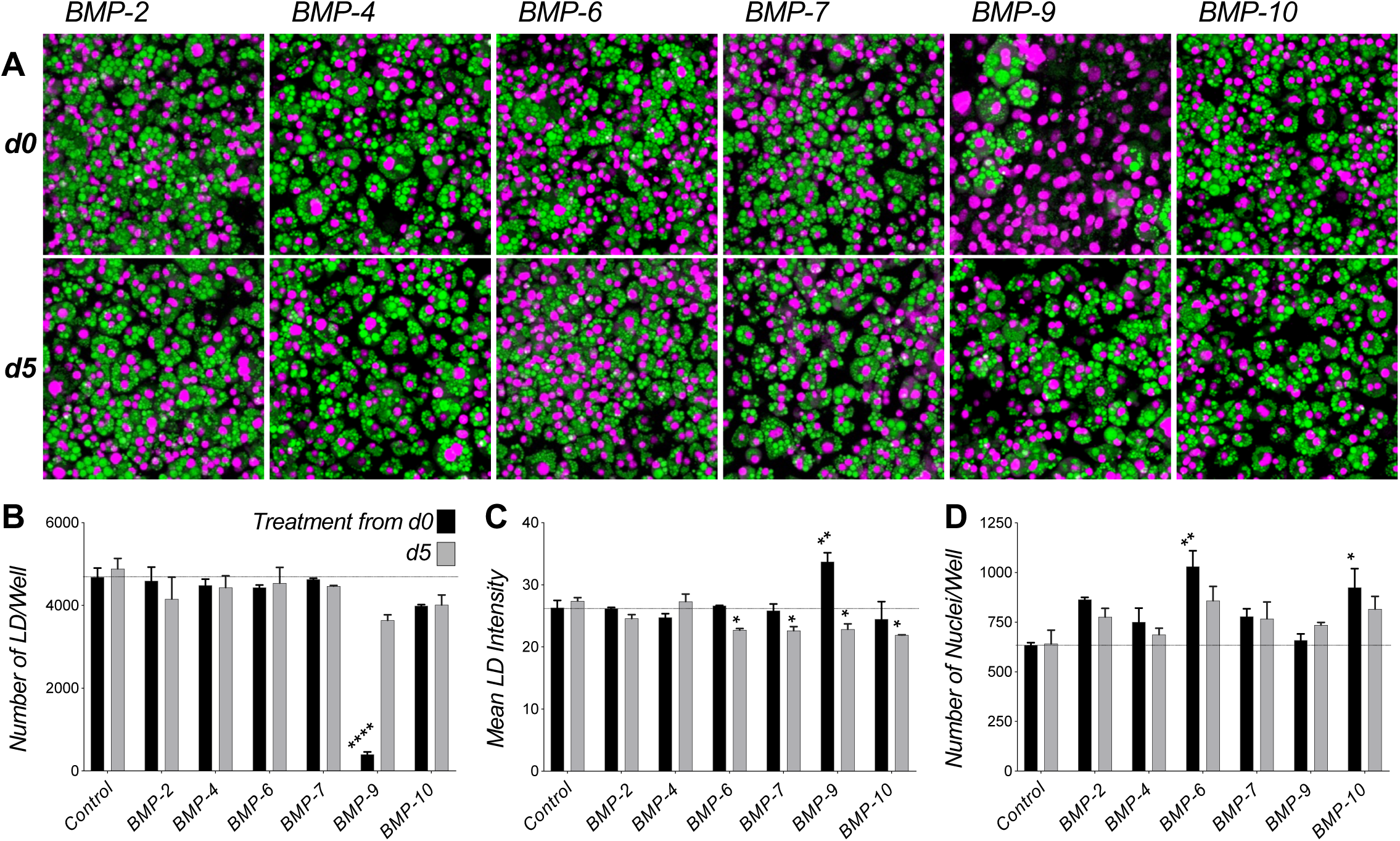
BMP-ligands induce 3T3-L1 proliferation. **A)** 3T3-L1 cells were grown with 1 nM ligand as noted or vehicle control (PBS) from day 0 (top panel) or day 5 (bottom panel) of differentiation until day 8. Vehicle control is shown in Figure 1B. Cells were fixed at day 8 and stained for lipids using Nile red (green), nuclei were counter-stained with DAPI (magenta). Cells treated with various BMPs generally show increased number of nuclei and DAPI fluorescence. **B-D)** Quantitative analysis of 3T3-L1 samples treated with ligands from day 0 (black) or day 5 (grey) of differentiation. Confocal images where analyzed using ImageJ and data were evaluated using GraphPad. Statistical significance from two biological replicates was calculated by two-way ANOVA and Dunnett’s multiple comparisons test (* p<0.05; **p<0.01; ***p<0.001, ****p<0.0001). **B)** Total number of lipid droplets (LD) per well. **C)** Mean lipid droplet intensity. **D)** Number of nuclei per well.

### Distinct expression of TGF-β family receptors and ligands in differentiating cells

To identify TGF-β family genes are expressed in undifferentiated 3T3-L1 cells that may contribute to adipogenic fate commitment, we analyzed their expression using publicly available microarray data (Figure S1) (Koster, Volckmann et al., 2019, Lattin, Schroder et al., 2008). We found several receptors were present at significant levels; however, the type I receptors ALK1, ALK6 and ALK7, as well as the type II receptors ActRIIB and AMHRII were only marginally expressed. Among co-receptors, only betaglycan (a.k.a. TGFβR3) was highly expressed, highlighting its recently described role in adipogenesis (Lee et al., 2019). The SMAD2/3 pathway activating ligands TGF-β1, TGF-β2, TGF-β3 and Activin A were also significantly expressed. This finding was surprising as exogeneous addition of these ligands blunted adipogenesis. However, expression of the gene *Inha* could potentially indicate how SMAD2/3 signaling is suppressed in these cells, as its gene product, Inhibin A, interacts with betaglycan to facilitate antagonism of Activin A (Lewis, Gray et al., 2000). Among SMAD1/5/8 pathway activating ligands, BMP-4 was most highly expressed, cementing its previously described role triggering adipocyte commitment from various precursors (Bowers & Lane, 2007, Tang et al., 2004). HUGO Gene names of TGFβ molecules are listed in Table S2 for reference.

### TGF-β family inhibitors prevent adipocyte precursor differentiation

To identify compounds that modulate adipogenic fates by inhibiting TGF-β family ligands, we investigated (in analogy to a high throughput drug screen) how 11 Fc-fusion traps that bind different groups of TGF-β family ligands affect 3T3-L1 differentiation. Traps were based on the ligand binding domains of TGF-β family type I receptors (ALK2, ALK3, ALK4), type II receptors (ActRIIA, ActRIIB, TGFβRII, BMPRII), antagonists (Cerberus) and co-receptors (mCryptic, Cripto-1, BAMBI) (Figures 3 and 4). Their ligand binding activities are summarized in Table S1 (Aykul & Martinez-Hackert, 2016a, Aykul & Martinez-Hackert, 2016b, Aykul, Parenti et al., 2017, Baud’huin, Solban et al., 2012, Sako, Grinberg et al., 2010). Of the tested traps, TGFβRII-Fc, mCryptic-Fc, and Cripto-1-Fc profoundly suppressed adipocyte formation from precursors, as indicated by the near total absence of LDs in cells treated at the beginning of differentiation (Figures 3 and 4). Matching our results with ligands, the few cells that differentiated in the presence of treatment had enlarged LDs (Figure 4B). Notably, most traps also reduced adipocyte proliferation (Figure 4 A,C). Thus, we have identified three Fc-fusion traps that prevent 3T3-L1 adipogenesis *in vitro*.

**Figure 3.**
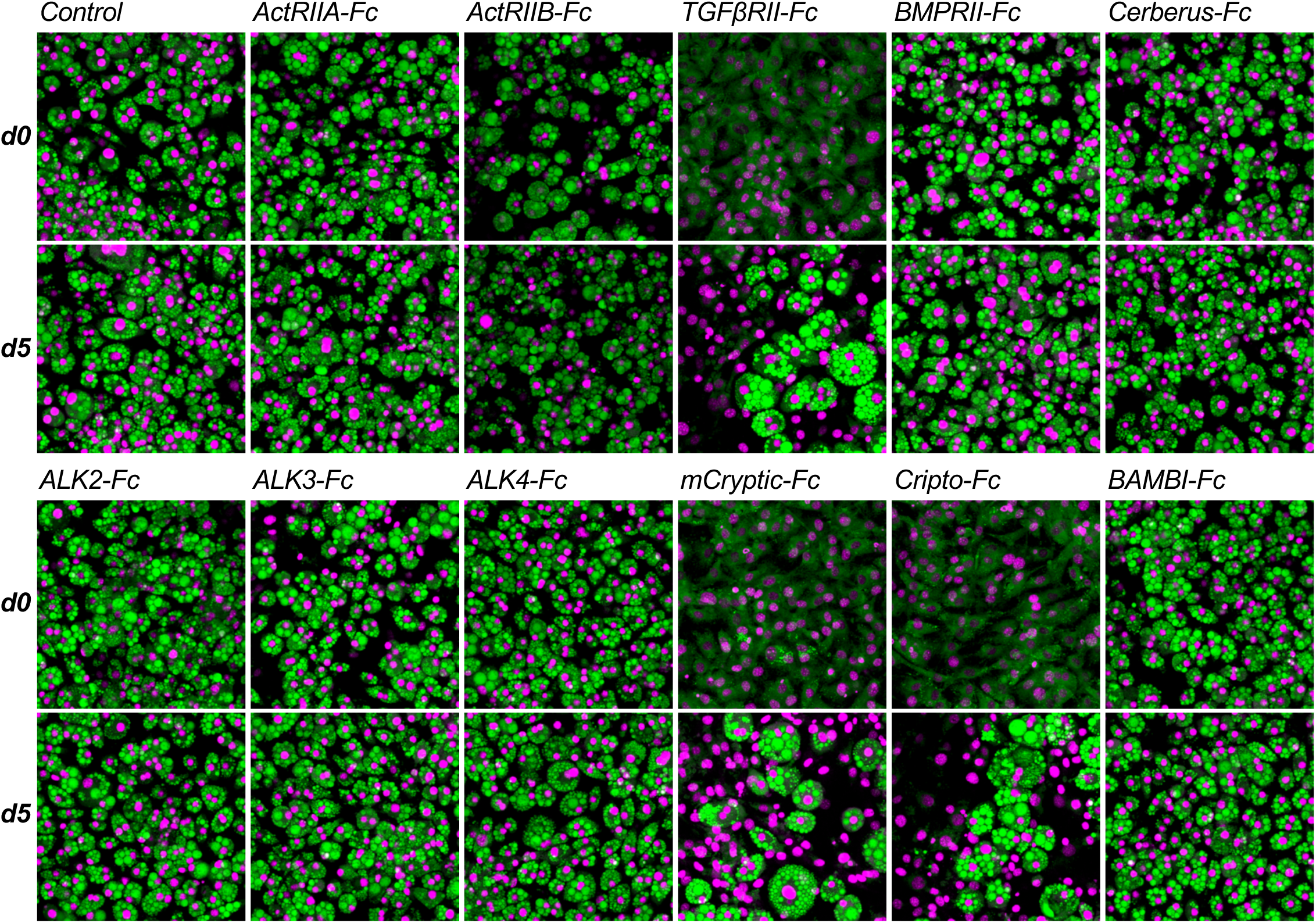
Differentiation assay screen identifies traps with anti-adipogenic activity. 3T3-L1 cells were grown in the presence of 300 nM Fc-fusion traps or vehicle control (PBS) as noted. Cells were treated from day 0 (top panel) or day 5 (bottom panel) of differentiation until day 8. Cells were fixed at day 8 and stained for lipids using Nile red (green), nuclei were counter-stained with DAPI (magenta). Cells treated with TGFβRII-Fc, mCryptic-Fc, or Cripto-1-Fc show significantly reduced lipid droplet formation. Each trap captures a unique group of ligands (ST1).

**Figure 4.**
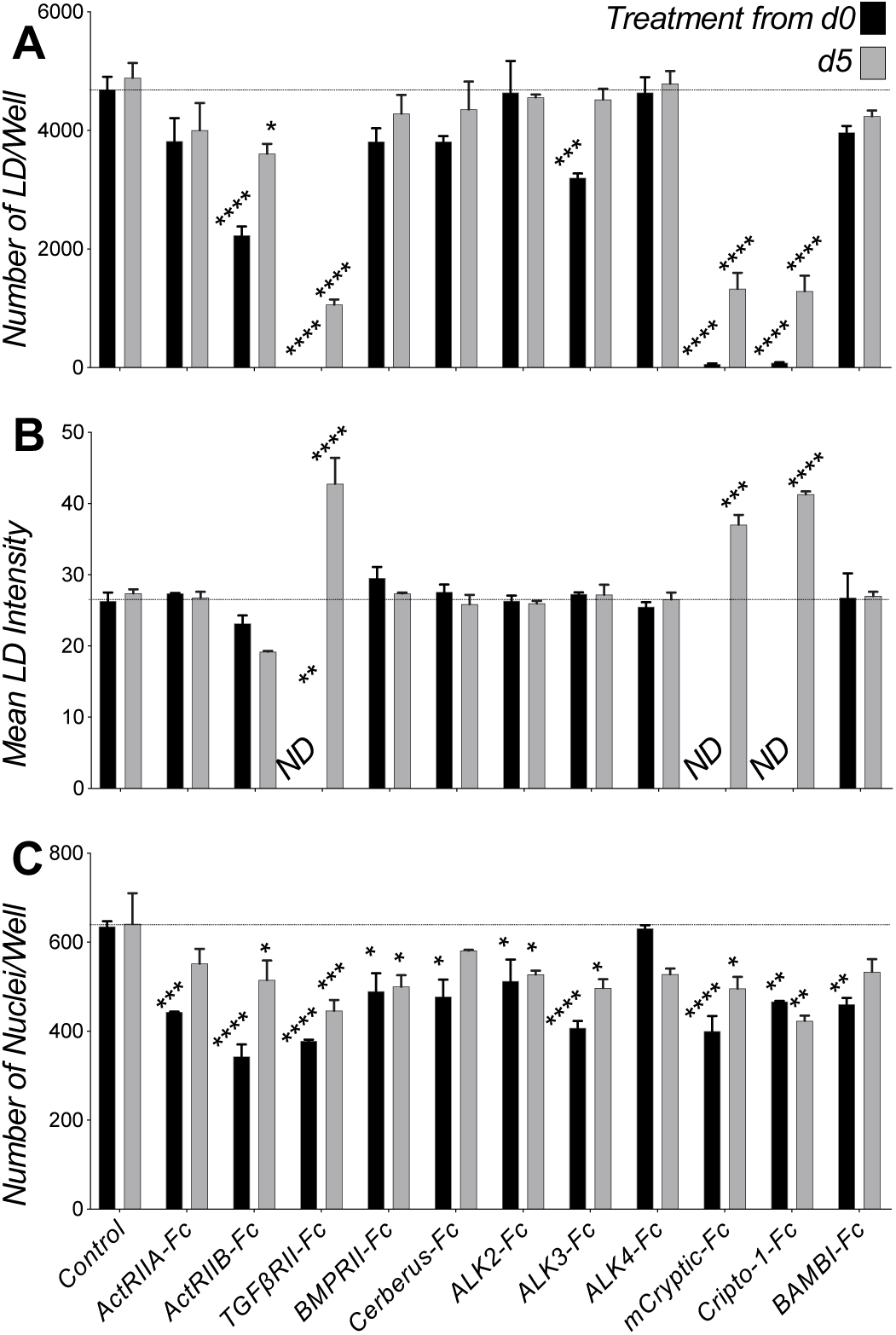
Inhibitory traps suppress lipid accumulation in differentiating 3T3-L1 cells. (**A-C)** Quantitative analysis of 3T3-L1 samples treated with ligands beginning at day 0 (black) or day 5 (grey) of differentiation. Confocal images where analyzed using ImageJ and GraphPad. Statistical significance from two biological replicates was calculated by two-way ANOVA and Dunnett’s multiple comparisons test (* p<0.05; **p<0.01; ***p<0.001, ****p<0.0001). **A)** Total number of lipid droplets (LD) per well. **B)** Mean lipid droplet intensity. **C)** Number of nuclei per well.

### Inhibitory traps suppress adipogenic transcription programs

To demonstrate that the inhibitory traps prevent adipogenesis, we investigated using qRT-PCR their effect on adipogenic gene expression (Figure 5) (Gjidoda, Tagore et al., 2014). In this experiment we focused on mCryptic-Fc and Cripto-1-Fc, as these are novel traps (Aykul et al., 2017). Both suppressed expression of adipogenic master regulators and other adipogenesis associated transcription factors, including peroxisome proliferator-activated receptor-γ and -δ (*Pparg* and *Ppard*) (Rosen, 2005). Although *Pparg* was induced 10.5-fold in untreated samples relative to undifferentiated controls, its expression was not induced in cells treated with mCryptic-Fc or Cripto-1-Fc (Figure 5A). Similarly, mRNA levels of the adipocyte lineage specific transcription factor C/EBPβ increased approximately 2.5-fold in differentiated control samples but were unchanged in treated samples (Figure 5B). In addition to adipogenic master regulators, both mCryptic-Fc and Cripto-1-Fc suppressed expression of adipocyte marker genes, including fatty acid binding protein 4 (*Fabp4*), cell death-inducing DFFA-like effectors a and c (*Cidea* and *Cidec*), perilipin (*Plin1*), adiponectin (*Adipoq*) and others (Figures 5C-H). For example, levels of the adipokine Adiponectin (*Adipoq*) increased about 400-fold in control samples but only 10-fold in treated samples (Figure 5C). Fatty acid-binding protein 4 (*Fabp4)* expression increased approximately 150-fold with differentiation in control samples but only about 2- to 4-fold in mCryptic-Fc or Cripto-1-Fc treated samples (Figure 5D). More strikingly, mRNA levels of *Cidec,* a regulator of adipocyte lipid metabolism that binds to lipid droplets and regulates their enlargement to restrict lipolysis and favor storage, increased over 5500-fold after differentiation in control samples but only 44- to 70-fold in treated samples (Figure 5E). Similarly, mRNA levels *Plin1*, a gene encoding for the lipid droplet-associated protein Perilipin, were increased approximately 970-fold in the control samples but levels in the mCryptic-Fc and Cripto-1-Fc treated samples only increased about 10-fold (Figure 5G). As we find that mCryptic-Fc and Cripto-1-Fc block expression of these adipogenic fate regulators and adipocyte marker genes (Hamza, Pott et al., 2009), we speculate that both traps, as well as TGFβRII-Fc and TGF-β-like ligands arrest adipocyte differentiation at a precursor stage or reprogram precursors to prevent adipogenic lineage commitment and/or adipogenesis.

**Figure 5.**
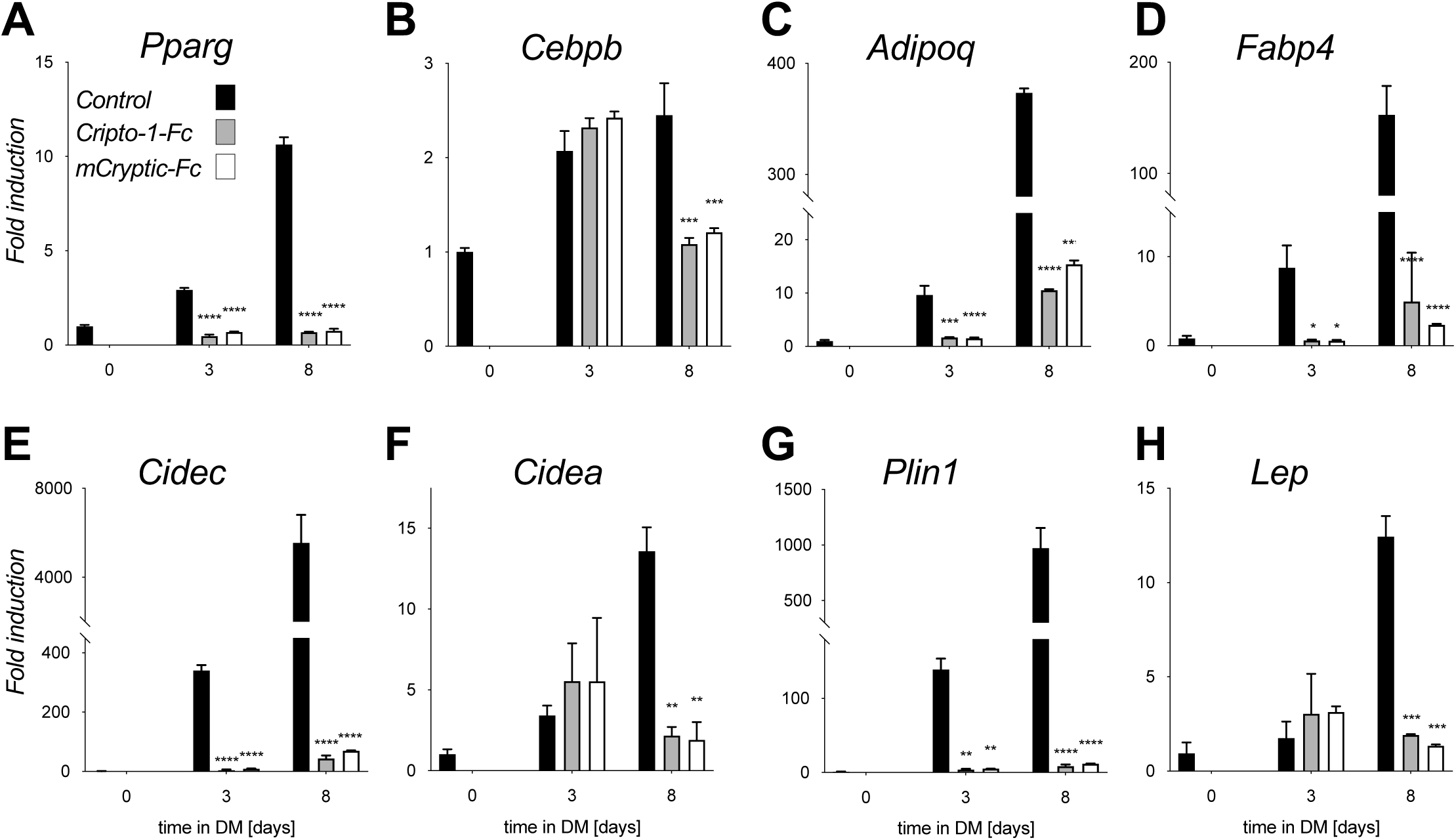
Inhibitory traps suppress expression of adipocyte marker genes. **A-H)** Induction of adipocyte marker gene expression was analyzed by qRT-PCR on days 0, 3 and 8 of differentiation in control, mCryptic-Fc, and Cripto-1-Fc treated cells (black, yellow and red bars, respectively). RNA isolation and qRT-PCR analysis were performed as described (Gjidoda et al., 2014). Data was normalized to Rpl4 mRNA and is shown as fold induction relative to day 0 levels. Statistical significance from two biological replicates was determined by two-way ANOVA and Sidaki’s post-hoc tests (* p<0.05; **p<0.01; ***p<0.001; ****p<0.0001) using GraphPad. Expression of adipogenic transcription factors **A)** *Pparg*, and **B)** *Cebpb.* Expression of adipocyte marker genes **C)** *Adipoq*, **D)** *Fabp4*, and **E)** *Lep*. Expression of genes associated with lipid droplet formation **F)** *Cidec*, **G)** *Cidea*, and **H)** *Plin1*. Expression of adipogenic genes is strongly induced by day 3 of differentiation. Cryptic-Fc and Cripto-1-Fc effectively suppress adipogenesis as evidenced by the absence of adipogenic gene expression.

### Distinct SMAD forms associate with adipose lineage commitment and inhibition

As SMAD transcription factors mediate intracellular responses to TGF-β family signals (Choy et al., 2000, Huang et al., 2009), we attempted to define their contribution in adipocyte development and adipogenesis inhibition. It is suggested that SMAD2/3 activation by TGF-βs, Activins or GDF-8 arrests adipocyte precursors in an undifferentiated state, while SMAD1/5/8 activation by BMPs and GDFs promotes adipocyte differentiation and hypertrophy (Choy et al., 2000, Derynck & Zhang, 2003, Huang et al., 2009, Schreiber et al., 2017, Shi & Massague, 2003). However, efforts to reveal the molecular basis of SMAD regulation have remained challenging (Choy et al., 2000), limiting our understanding of a key mechanism associated with AT development.

To define the functional roles of endogenous SMADs, we probed their activation by Western blot (Figure 6A-C). Microarray data indicate that most SMADs are expressed in undifferentiated 3T3-L1 cells, except for the weakly expressed SMAD7 and SMAD8 (Figure S1) (Koster et al., 2019, Lattin et al., 2008). Using a p-SMAD1/5/8 monoclonal antibody, we observed a strong 50 kDa band in 3T3-L1 cells at the beginning of differentiation (day 0), indicating that SMAD1/5/8 are C-terminally phosphorylated (*i.e.* activated) in adipocyte precursors (Figure 6A, upper panel). The 50 kDa p-SMAD1/5/8 signal became considerably weaker after day 5 but persisted in all samples that differentiated (Figures 6A,B). Strikingly, the 50 kDa p-SMAD1/5/8 signal was almost completely superseded by a new, higher molecular weight species of 55-60 kDa in all cells treated with traps or ligands that arrested differentiation (Figures 6A,B, upper panel), whereas traps that did not affect differentiation retained the lower molecular weight species. We refer hereafter to the 50 and 55-60 kDa bands as p-SMAD1/5^Lo^ and p-SMAD1/5^Hi^, respectively.

**Figure 6.**
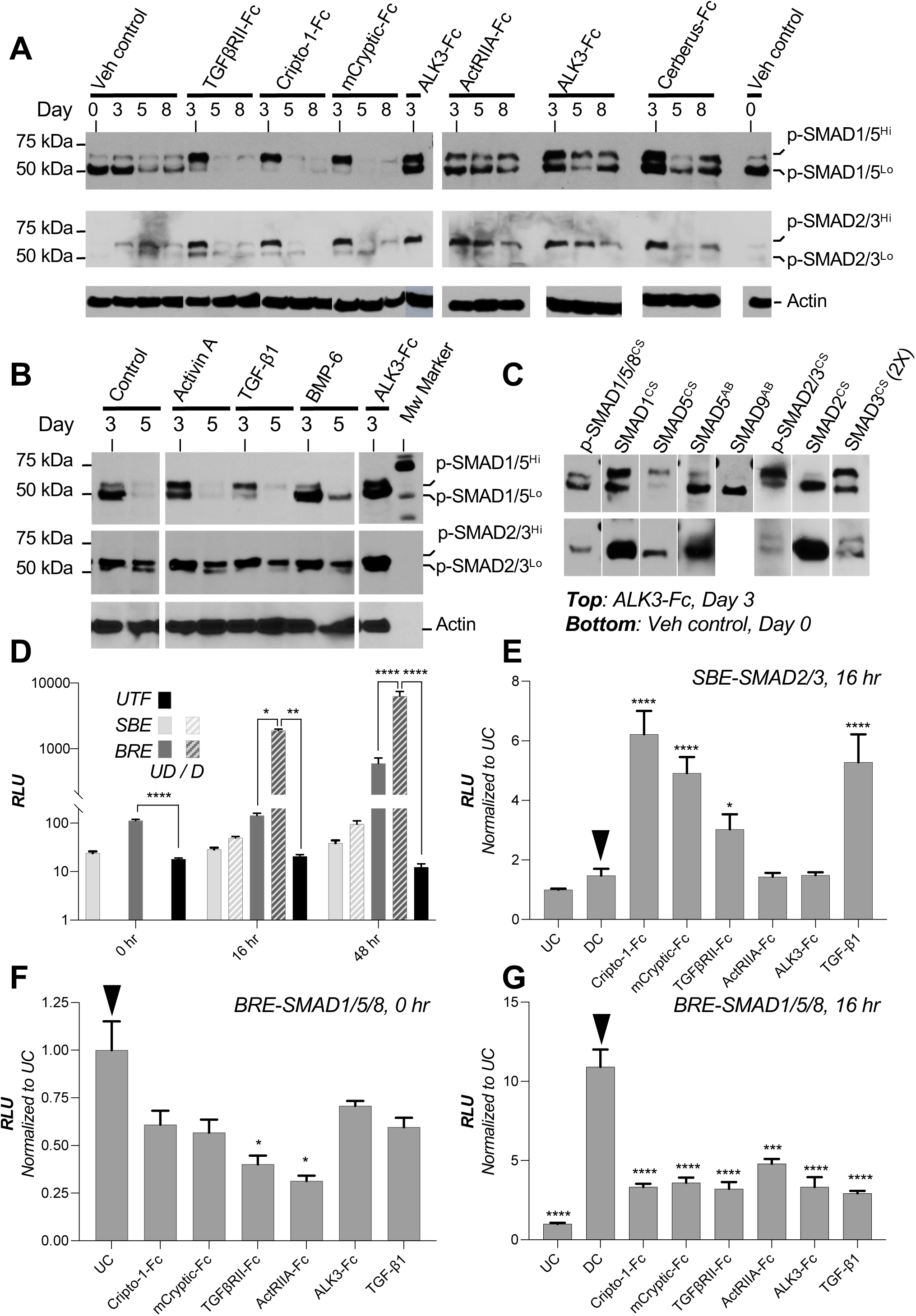
Electrophoretic mobility shift in p-SMAD1/5 and altered SMAD activities associate with anti-adipogenic mechanism. **A)** Anti-p-SMAD Western blots of whole cell lysate show C-terminal Serine phosphorylation of SMADs (top panel: p-SMAD1/5/8, middle panel: p-SMAD2/3) in samples treated with different ligand traps or vehicle (PBS). Treatments were started at day 0 of differentiation and samples were collected at days 0, 3, 5, or 8 of differentiation as noted. Two p-SMAD1/5/8 forms were detected and are labeled p-SMAD1/5^Lo^ and p-SMAD1/5^Hi^, reflecting differences in electrophoretic mobility. Similarly, p-SMAD2/3 also appears as two bands. Actin Western blots were used as sample loading controls. Blots were loaded with 10 µg protein per lane. **B)** Anti-p-SMAD Western blots of whole cell lysate show C-terminal Serine phosphorylation of SMADs (top panel: p-SMAD1/5/8, middle panel: p-SMAD2/3) in samples treated with different ligands and Fc control (ALK3-Fc). Treatments were started at day 0 of differentiation. Samples were loaded at a 3-fold higher amount than in panel A to make up lower detection or lower abundance issues. Samples were collected at days 3 or 5 of differentiation. Actin was used as loading control. Blots for Actin and p-SMAD1/5/8 have 10 µg protein per lane. The p-SMAD2/3 blot has 30 µg protein per lane. **C)** Anti-SMAD Western blots of whole cell lysate show overall SMAD levels in ALK3-Fc treated samples (top panel, sample collected at day 3) and vehicle control samples (PBS, bottom panel, sample collected at day 0). ALK3-Fc was used as both p-SMAD1/5^Lo^ and p-SMAD1/5^Hi^ forms were prominently visible with his treatment. Antibodies used are noted above the blots. CS denotes antibody from Cell Signaling Technologies, AB denotes antibody from Abcam. p-SMAD antibodies recognize conserved, C-terminally phosphorylated Serine residues in activated SMADs. All other SMAD antibodies were raised against unique sequences within each SMAD protein. Blots were loaded with 10 µg protein per lane. **D)** Dual luciferase reporter assay in 3T3-L1 cells shows basal SMAD1/5/8 and SMAD2/3 signaling as noted at 0 (light grey) and 16 (dark grey) hours of differentiation. Samples represented by solid bars were undifferentiated (UD), samples represented by stripped bars were treated with differentiation reagents (D). Firefly luciferase reporter lacking control (untransfected, UTF) is shown as black bar. Data were analyzed using GraphPad and statistical significance from four biological replicates was determined by one-way ANOVA and Fisher’s LSD tests (* p<0.05; **p<0.01; ***p<0.001). **E)** Dual luciferase reporter assay in 3T3-L1 cells shows SMAD2/3 signaling at 16 h of differentiation. Samples were treated with different ligands or traps as noted. Both reporter plasmid transfection and treatment started at 0 hours of differentiation. The SMAD2/3 mediated firefly luciferase signal was normalized against the *Renilla* luciferase internal control. Data are shown as fold induction relative to undifferentiated control (UC). Data were analyzed using GraphPad. Statistical significance from four biological replicates was determined by one-way ANOVA and Fisher’s LSD tests by comparing treatments against differentiated control (DC, black arrow). **F)** Dual luciferase reporter assay in 3T3-L1 cells shows SMAD1/5/8 signaling at 0 h of differentiation. Samples were treated with different ligands or traps. Both reporter plasmid transfection and treatment started at day (−1). The SMAD1/5/8 dependent firefly luciferase signal was measured at 0 h of differentiation and normalized against *Renilla* luciferase control. Data are shown as fold induction relative to undifferentiated control (UC). Statistical significance from four biological replicates was determined by one-way ANOVA and Fisher’s LSD tests by comparing treatments against undifferentiated control (UC, black arrow). **G)** Dual luciferase reporter assay in 3T3-L1 cells shows SMAD1/5/8 signaling at 16 h of differentiation. Samples were treated with different ligands or traps as noted. Both reporter plasmid transfection and treatment started at 0 hours of differentiation. The SMAD1/5/8 dependent firefly luciferase signal was normalized against *Renilla* luciferase control. Data are shown as fold induction relative to undifferentiated control (UC). Data were analyzed using GraphPad. Statistical significance from four biological replicates was determined by one-way ANOVA and Fisher’s LSD tests by comparing treatments against differentiated control (DC, black arrow, * p<0.05; **p<0.01; ***p<0.001, ***p<0.0001).

To establish the identity of the two bands, we used antibodies against unique sequences within the SMAD linker regions (Figure 6C). Both p-SMAD1/5^Lo^ and p-SMAD1/5^Hi^ bands reacted with a SMAD1 specific antibody. In addition, at least one anti-SMAD5 and one anti-SMAD8 antibody reacted with the band corresponding to p-SMAD1/5^Lo^. However, neither anti-SMAD5 nor anti-SMAD8 antibodies reacted significantly with the p-SMAD1/5^Hi^ form. These results are consistent with earlier knock-down data identifying SMAD1 and SMAD5 forms of distinct molecular weights (Daly, Randall et al., 2008) and suggest that SMAD1 is the major SMAD associated with the p-SMAD1/5^Hi^ species. In contrast to SMAD1/5/8, we only observed weak 50 and 55-60 kDa bands with the p-SMAD2/3 antibody in untreated cells at days 3 and 5 of differentiation at the same gel loading concentrations (Figure 6A, middle panel). Notably, cells treated with either inhibitory or non-inhibitory traps or ligands, or with *bona fide* SMAD2/3 activation inhibitors, including TGFβRII-Fc and ActRIIA-Fc, presented a stronger p-SMAD2/3 Western blot signal at day 3 of differentiation (Figure 6A,B, middle panel). Based on these observations, we propose that the key signal associated with 3T3-L1 commitment and differentiation is p-SMAD1/5^Lo^. Adipogenesis inhibitors did not alter C-terminal SMAD1/5/8 phosphorylation but induced a near complete transition to the p-SMAD1/5^Hi^ form, which was fundamentally associated with differentiation arrest. By contrast, SMAD2/3 was minimally activated in these cells and activation as indicated by C-terminal SMAD2/3 phosphorylation did not appear to affect 3T3-L1 differentiation.

### Reduced SMAD1/5/8 and increased SMAD2/3 signaling associate with adipogenesis arrest

To elucidate the functional consequences of the p-SMAD1/5^Lo^ to p-SMAD1/5^Hi^ transition, we examined SMAD1/5/8 signaling activities using the SMAD1/5/8 responsive BRE reporter (von Bubnoff, Peiffer et al., 2005). Looking at basal levels, we observed a persistent SMAD1/5/8 mediated luciferase signal that was 6- to 7-fold greater relative to un-transfected cells in undifferentiated control cells at 0- and 16-hours post confluence (Figure 6D). The signal increased significantly in cells treated with differentiation medium relative to the undifferentiated control at 16- and 48-hours post differentiation (Figure 6D). Significantly, cells treated with adipogenesis inhibitors had 4- to 6-fold reduced SMAD1/5/8 signaling relative to untreated controls (Figure 6F,G). Thus, consistent with our findings of C-terminally phosphorylated p-SMAD1/5^Lo^ (Figure 6A), SMAD1/5/8 signaling is activated in 3T3-L1 cells and increases further with differentiation. Adipogenesis inhibitors suppressed the SMAD1/5/8 signal, even as the p-SMAD1/5^Hi^ C-terminus remained phosphorylated (Figure 6A), indicating that p-SMAD1/5^Hi^ loses the ability to activate BRE mediated transcription.

In contrast to SMAD1/5/8, basal SMAD2/3 signaling activity as measured by the SMAD2/3 responsive SBE reporter was low and minimally induced upon differentiation (Figure 6D). Strikingly, all adipogenesis inhibitors, including Cripto-1-Fc, TGFβRII-Fc and TGF-β1, increased the SMAD2/3 dependent reporter signal approximately 3- to 6-fold relative to untreated controls (Figure 6E). These findings agree with our Western blot results showing induction of the p-SMAD2/3 form by these treatments (Figure 6B). However, in contrast to our Western blot findings, non-inhibitory traps did not induce SMAD2/3 dependent luciferase expression (Figure 6A,B), revealing a functional distinction between C-terminal SMAD2/3 phosphorylation and SBE dependent transcriptional activity. Nevertheless, these findings support the conclusion that both activation of SMAD2/3 and inhibition of SMAD1/5/8 signaling via induction of the p-SMAD1/5^Hi^ form leads to adipogenesis arrest.

### p-SMAD1/5^Hi^ is hyper-phosphorylated and does not translocate to the nucleus

Although C-terminal SMAD Serine phosphorylation is the key step in TGF-β family receptor mediated SMAD pathway activation (Derynck & Zhang, 2003, Shi & Massague, 2003), Serine/Threonine kinase dependent SMAD hyper-phosphorylation is also known to regulate SMAD activities (Alarcon, Zaromytidou et al., 2009, Aubin, Davy et al., 2004, Fuentealba, Eivers et al., 2007, Kamato, Burch et al., 2013, Kretzschmar, Doody et al., 1997, Kuroda, Fuentealba et al., 2005, Moustakas, Souchelnytskyi et al., 2001, Pera, Ikeda et al., 2003, Sapkota, Alarcon et al., 2007). As we detected an increase in apparent p-SMAD1/5 molecular weight following treatment with adipogenesis inhibitors, we hypothesized that the p-SMAD1/5^Hi^ species could represent a hyper-phosphorylated form (Alarcon et al., 2009, Aubin et al., 2004, Fuentealba et al., 2007, Kamato et al., 2013, Kretzschmar et al., 1997, Kuroda et al., 2005, Pera et al., 2003, Sapkota et al., 2007).

To test this hypothesis, we first treated 3T3-L1 lysates of control and TGFβRII-Fc treated samples with alkaline phosphatase (AP) and evaluated changes in SMAD1/5 electrophoretic mobility using different antibodies (Figure 7A). As expected, AP treated samples were largely undetectable by the p-SMAD1/5/8 antibody due to near complete loss of all phosphate groups (Figure 7A, left panel). By contrast, SMAD1 and SMAD5 antibodies reacted well with both untreated and AP treated samples (Figure 7A). The difference in electrophoretic mobility following AP digestion was small in control samples (C), indicating that the p-SMAD1/5^Lo^ form is phosphorylated at a few sites, including its C-terminus. However, the difference in electrophoretic mobility following AP digestion in TGFβRII-Fc (T-Fc) treated samples was much greater, indicating that the p-SMAD1/5^Hi^ form is phosphorylated at additional sites (Figure 7A, middle and right-side panels). Based on this evidence, we propose that hyper-phosphorylation effects the shift in electrophoretic mobility between the p-SMAD1/5^Lo^ to p-SMAD1/5^Hi^ forms.

**Figure 7.**
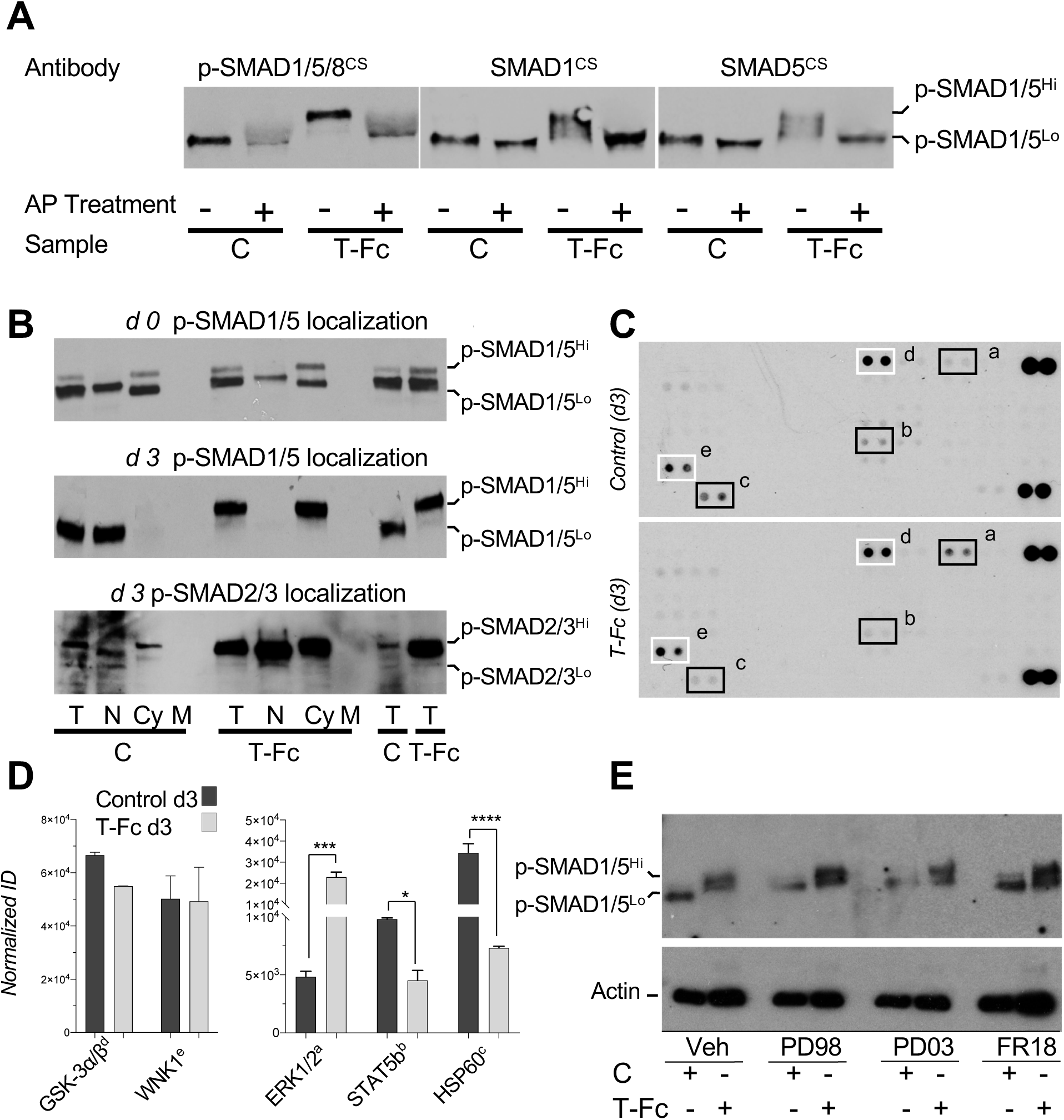
Electrophoretic mobility shift is associated with p-SMAD1/5 hyper-phosphorylation and leads to reduced p-SMAD1/5 nuclear translocation. **A)** Analysis of SMAD1/5/8 linker phosphorylation by Alkaline phosphatase digestion. Western blots of whole cell lysate show Vehicle control (C) and TGFβRII-Fc (T-Fc) treated samples subjected to Alkaline phosphatase (AP) dephosphorylation (+). Antibodies against C-terminally phosphorylated SMAD1/5/8 (p-SMAD1/5/8), SMAD1 and SMAD5 were used as noted. The p-SMAD1/5^Hi^ form found in TGFβRII-Fc treated samples reduces to the p-SMAD1/5^Lo^ form upon AP digestion. **B)** SMAD localization by cellular fractionation. Total cell lysate (T), nuclear (N), cytoplasmic (Cy), and membrane (M) fractions of vehicle control (C) and TGFβRII-Fc (T-Fc) treated samples were analyzed by Western blot after cellular fractionation using p-SMAD1/5/8 and p-SMAD2/3 antibodies. The top panel and the middle and bottom panels, respectively, show samples with treatment beginning at d-2 and d0 and collected at d0 and d3 of differentiation. p-SMAD1/5^Hi^ is found in the cytoplasmic fraction in all TGFβRII-Fc treated samples. Only p-SMAD1/5^Lo^ is found in the nuclear fraction. **C)** Membrane-based sandwich immunoassay analysis of kinase phosphorylation (RnD Systems Proteome Profiler Human Phospho-Kinase Array Kit, ARY003B) in Vehicle Control and TGFβRII-Fc treated samples collected at day 3 of differentiation. Boxed immunoblots show statistically significantly differences. Black boxes represent proteins that are differentially phosphorylated between Control (C) and TGFβRII-Fc (T-Fc) treated samples. White boxes represent proteins that are phosphorylated in all 3T3-L1 samples. Box (a) corresponds to ERK1/2, box (b) to STAT5b, box (c) to HSP60, box (d) to GSK-3α/β and box (e) to WNK1. **D)** Quantitative analysis of Human Phospho-Kinase Array blots. Spot integrated densities were determined with ImageJ. Intensities were normalized against capture control antibody spots. Statistical significance from two biological replicates was determined by one-way ANOVA and Fisher’s LSD tests (* p<0.05; **p<0.01; ***p<0.001) using GraphPad. **E)** Effect of ERK inhibitors on SMAD1/5/8 hyper-phosphorylation. Control (C) and TGFβRII-Fc (T-Fc) treated samples 3T3-L1 cells were simultaneously treated with vehicle (Veh, DMSO) or with the ERK inhibitors PD98059 (PD98, 1 μM), PD0325901 (PD03, 1 μM), and FR180204 (FR18, 10 μM). Equal amounts (10 μg) of protein from 3T3-L1 cells collected at day 3 of differentiation were analyzed by Western blot using a specific p-SMAD1/5/8 antibody. Actin loading controls are also shown (bottom panel). ERK inhibitors do not prevent formation of the p-SMAD1/5^Hi^ form.

To determine if hyper-phosphorylation alters cellular localization of p-SMAD1/5^Hi^ as previously indicated (Kretzschmar et al., 1997, Kuroda et al., 2005, Pera et al., 2003, Sapkota et al., 2007), we separated cell extracts into nuclear, cytoplasmic and membrane fractions and probed these with anti-p-SMAD antibodies (Figure 7B). The p-SMAD1/5^Lo^ form was detectable in both cytosolic and nuclear fractions, whereas p-SMAD1/5^Hi^ was only detectable in the cytoplasmic fraction (Figure 7B, top and middle panels). Untreated control cells presented both p-SMAD1/5 forms at the beginning of differentiation and mainly exhibited higher levels of nuclear pSMAD1/5^Lo^ during differentiation. This finding is consistent with the observed induction of the BRE reporter activity. By contrast, cells treated with the adipogenesis inhibitor TGFβRII-Fc only exhibited the p-SMAD1/5^Hi^ form in the cytoplasmic fraction (Figure 7B), a finding that is consistent with reduced signaling. Collectively, these and reporter assay results shown in Figure 6 indicate that the p-SMAD1/5^Lo^ form can translocate to the nucleus and activate transcription, whereas p-SMAD1/5^Hi^ remains cytoplasmic and exhibits significantly reduced transcriptional activity. Notably, we did not observe differences in localization of p-SMAD2/3 during differentiation between control and inhibitor treated samples (Figure 7B, bottom panel). However, consistent with Figure 6, we found p-SMAD2/3 levels increased significantly in TGFβRII-Fc treated samples.

### Kinase activation and SMAD1/5/8 hyper-phosphorylation

To identify pathways that lead to SMAD1/5/8 hyper-phosphorylation, we investigated intracellular serine/threonine kinase activation using the phospho-kinase profiler array kit (Figure 7C, D). We focused on this group of enzymes as several kinases, including ERK, JNK and PI3K, are thought to hyper-phosphorylate SMADs at their interdomain linker region and thus regulate SMAD activities (Alarcon et al., 2009, Aubin et al., 2004, Fuentealba et al., 2007, Kamato et al., 2013, Kretzschmar et al., 1997, Kuroda et al., 2005, Pera et al., 2003, Sapkota et al., 2007). All tested samples, including undifferentiated precursors and differentiating adipocytes, showed significant GSK-3α/β and WNK1 phosphorylation, indicating that these kinases may potentially have relevant roles in adipogenic lineage commitment but may not be involved in differentiation arrest. Notably, ERK phosphorylation was increased, while phosphorylation of STAT5b and the heat shock protein HSP60 was decreased in cells treated with adipogenesis inhibitors, further indicating that the three proteins have roles in adipogenesis (Bost, Aouadi et al., 2005, Gulden, Mollerus et al., 2008, Stephens, Morrison et al., 1999). As ERK activation has been linked with SMAD hyper-phosphorylation (Hayashida, Decaestecker et al., 2003, Hough, Radu et al., 2012, Matsuura, Wang et al., 2005), we speculated that ERK inhibitors could potentially suppress formation of the p-SMAD1/5^Hi^ species. However, all tested ERK inhibitors, including the ERK phosphorylation inhibitors PD98059 and PD0325901, and the selective ERK inhibitor FR180204, failed to prevent SMAD1/5/8 hyper-phosphorylation or rescue adipogenesis at their highest non-toxic concentrations, indicating that ERK kinase may not directly give rise to the p-SMAD1/5^Hi^ species in this context (Figure 7E).

### Small molecule inhibitors reveal functional crosstalk between SMAD2/3 and SMAD1/5/8 pathways

As our Western blot and reporter assays indicated that both reduced SMAD1/5/8 and increased SMAD2/3 signaling resulted in adipogenesis inhibition, we wanted to determine if these mechanisms are independent or the two SMAD branches interact to regulate adipocyte development. To elucidate the mechanistic contributions of each SMAD branch and identify a functional interplay, we examined the effects of the SMAD2/3 and SMAD1/5/8 activation inhibitors SB-431542 (SB43) and LDN-193189 (LDN) on adipocyte differentiation at their highest non-toxic concentrations (Figure 8) (Cuny, Yu et al., 2008, Inman, Nicolas et al., 2002). Although SB43 alone did not have a detectable effect on adipogenesis, it fully rescued adipocyte formation in cells treated with inhibitors (Figure 8A, top and middle panel), indicating that SMAD2/3 activation arrests precursors in an undifferentiated state or reprograms 3T3-L1 cells toward a non-adipogenic lineage. However, LDN alone also inhibited adipogenesis as evidenced by the complete lack of lipid droplet in LDN treated cells (Figure 8A, top and bottom panel). This observation is consistent with previous findings (Suenaga, Kurosawa et al., 2013) and provides direct evidence that SMAD1/5/8 signaling is required for adipogenesis.

**Figure 8.**
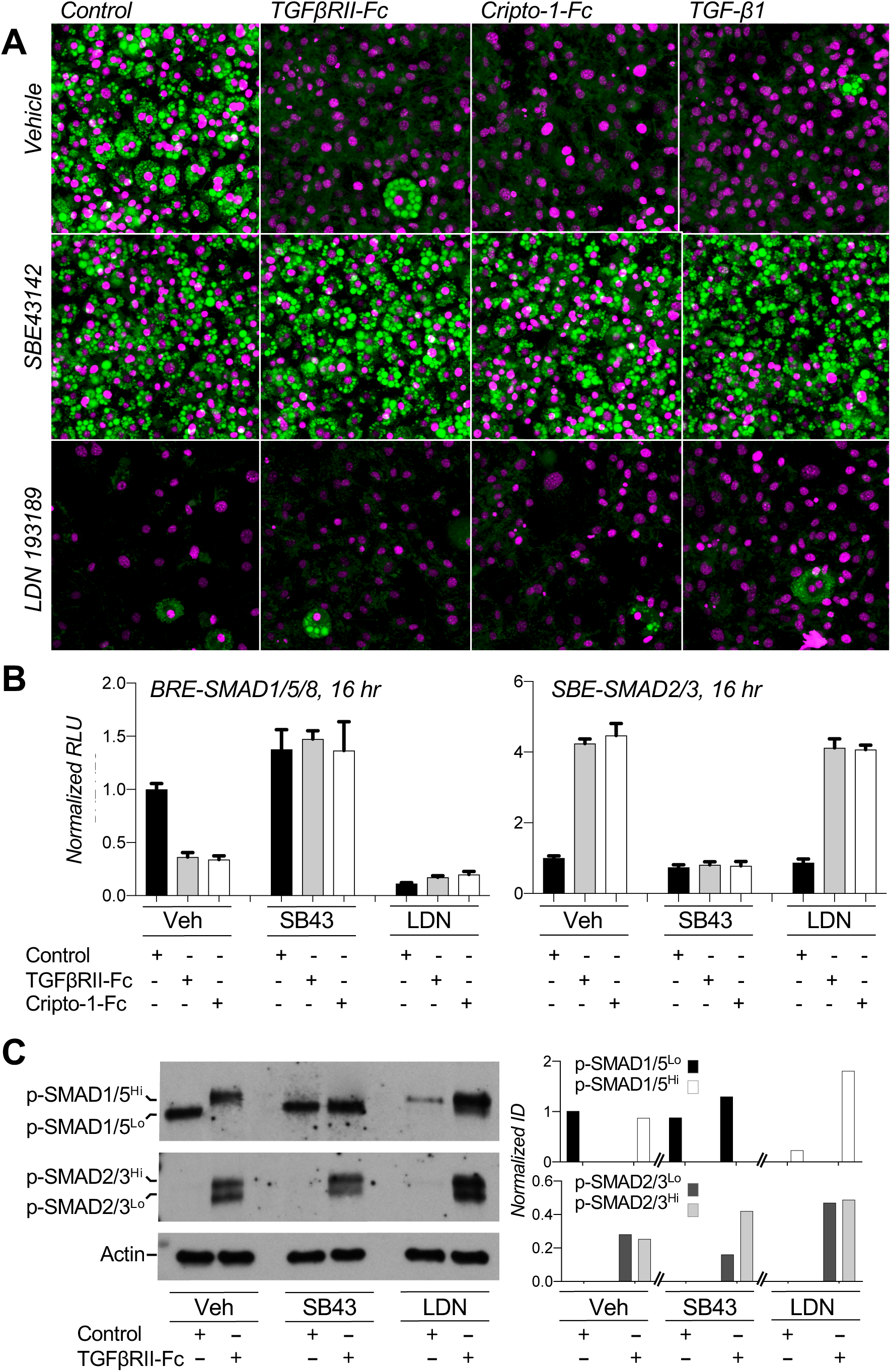
Effect of small molecule inhibitors on 3T3-L1 differentiation and signaling. **A)** 3T3-L1 cells were grown in the presence of 300 nM Fc-fusion traps or vehicle control (PBS) and the small molecule inhibitors SB-43142 (SB43, 10 μM) or LDN 193189 (LDN, 1 μM) as noted. Cells were treated from day 0 of differentiation until collected. Cells were collected at day 8, fixed and stained for lipids using Nile red (green). Nuclei were counter-stained with DAPI (magenta). SB43 rescues cells treated with adipogenesis inhibitors, while LDN suppresses adipogenesis as indicated by the respective presence and absence of lipid droplets (green). **B)** Dual luciferase reporter assay shows BRE-SMAD1/5/8 (left panel) and SBE-SMAD2/3 (right panel) mediated signaling in 3T3-L1 cells at 16 hours of differentiation. Black bars correspond to vehicle, light grey bars to TGFβRII-Fc, white bars to Cripto-1-Fc treatment. Groups are separated by secondary treatment as noted, including Veh (DMSO), SB43 and LDN. SMAD2/3 or SMAD1/5/8 dependent firefly luciferase signals were normalized against their respective *Renilla* luciferase internal controls. All treatments were then normalized to Vehicle controls (black bar of corresponding Veh group). Controls in vehicle group were used as reference for analysis in GraphPad. Statistical significance from four biological replicates was determined by one-way ANOVA and Fisher’s LSD tests by comparing treatments against differentiated control (DC). black arrow, * p<0.05; **p<0.01; ***p<0.001, ***p<0.0001). SB43 induces BRE-SMAD1/5/8 luciferase activity. LDN does not affect SBE-SMAD2/3 luciferase activity. **C)** Anti-p-SMAD Western blots of 3T3-L1 cells treated with small molecule inhibitors. The top panel shows p-SMAD1/5/8, the middle panel p-SMAD2/3, the bottom panel actin loading controls. Control and TGFβRII-Fc treated cells were subjected to additional Veh (DMSO), SB43 and LDN treatment from day 0 of differentiation and samples were collected at day 3. SB43 prevents SMAD1/5/8 hyper-phosphorylated in TGFβRII-Fc treated cells as evidenced by the absence of pSMAD1/5^Hi^.

To establish how SB43 and LDN alter SMAD signaling and elicit their respective effects, we investigated their activities using SMAD responsive luciferase reporter assays (Figure 8B). As shown in Figure 6, BRE-SMAD1/5/8 dependent luciferase expression was significantly attenuated by the adipogenesis inhibitor traps TGFβRII-Fc and Cripto-1-Fc, and by TGF-β1 (Figure 8B, Veh). Strikingly, SB43 induced SMAD1/5/8 signaling in control cells and rescued SMAD1/5/8 signaling in cells treated with inhibitor traps (Figure 8B, SB43), indicating that SMAD2/3 activation or the SB43 target kinases ALK4, ALK5 or ALK7 directly or indirectly regulate SMAD1/5/8 activities in these cells. In contrast to SB43, LDN suppressed SMAD1/5/8 signaling (Figure 8B, LDN). This finding is consistent with its mechanism of action and provides further evidence that SMAD1/5/8 activation is essential for 3T3-L1 adipogenesis. Remarkably, TGFβRII-Fc and Cripto-1-Fc induced SBE-SMAD2/3 dependent luciferase expression (Figures 6E and 8B, Veh). This activation was inhibited by SB43 but not LDN as expected (Figure 8B), indicating that SMAD2/3 activation is independent of SMAD1/5/8. Thus, using SMAD2/3 and SMAD1/5/8 pathway inhibitors, we showed that suppressed SMAD2/3 and activated SMAD1/5/8 signaling are critical for 3T3-L1 adipogenesis. Notably, SMAD2/3 activation was coupled to SMAD1/5/8 inhibition. SB43 could, therefore, rescue SMAD1/5/8 dependent reporter activity and adipogenesis in cells treated with TGFβRII-Fc and Cripto-1-Fc.

To determine at a molecular level how SB43 and LDN elicit their effects, we probed SMAD activation states using p-SMAD antibodies (Figure 8C). As we showed in Figure 6, control samples only exhibited the p-SMAD1/5^Lo^ species, while cells treated with TGFβRII-Fc only exhibited p-SMAD1/5^Hi^ (Figure 8C, top panel). Strikingly, SB43 suppressed formation of the p-SMAD1/5^Hi^ form in TGFβRII-Fc treated 3T3-L1 cells, indicating that pathways or targets inhibited by SB43 could be directly linked with SMAD1/5/8 hyper-phosphorylation. Although LDN suppressed C-terminal SMAD1/5/8 phosphorylation in control cells as expected (Figure 8C), it did not prevent SMAD1/5/8 C-terminal or hyper-phosphorylation in cells co-treated with TGFβRII-Fc. Together with reporter assay data (Figure 8B), these data therefore suggest that the p-SMAD1/5^Hi^ species does not activate transcription of canonical p-SMAD1/5/8 target genes in differentiating 3T3-L1 cells.

In contrast to SMAD1/5/8, p-SMAD2/3 was induced in all cells treated with TGFβRII-Fc, a surprising finding that was nevertheless consistent with luciferase reporter assays (Figure 8C, middle panel). Strikingly, although SB43 inhibited SMAD2/3 signaling in TGFβRII-Fc treated cells, it did not reduce SMAD2/3 C-terminal phosphorylation, as evidenced by the p-SMAD2/3 specific immunoblot. However, SMAD2/3 presented mainly as a p-SMAD2/3^Hi^ form, suggesting that differential activation of SMAD2 vs. SMAD3 may effect the distinct signaling outcomes (Nakao, Imamura et al., 1997). In conclusion, using small molecule inhibitors we show that SMAD1/5/8 signaling is essential whereas SMAD2/3 signaling is detrimental for 3T3-L1 adipogenesis (Figure 9). SMAD1/5/8 hyper-phosphorylation, which consistently associates with reduced SMAD1/5/8 signaling and adipogenesis arrest, appears to be directly regulated by SMAD2/3 activation or by SB43 target kinases, potentially revealing a functional crosstalk between the two SMAD branches.

**Figure 9.**
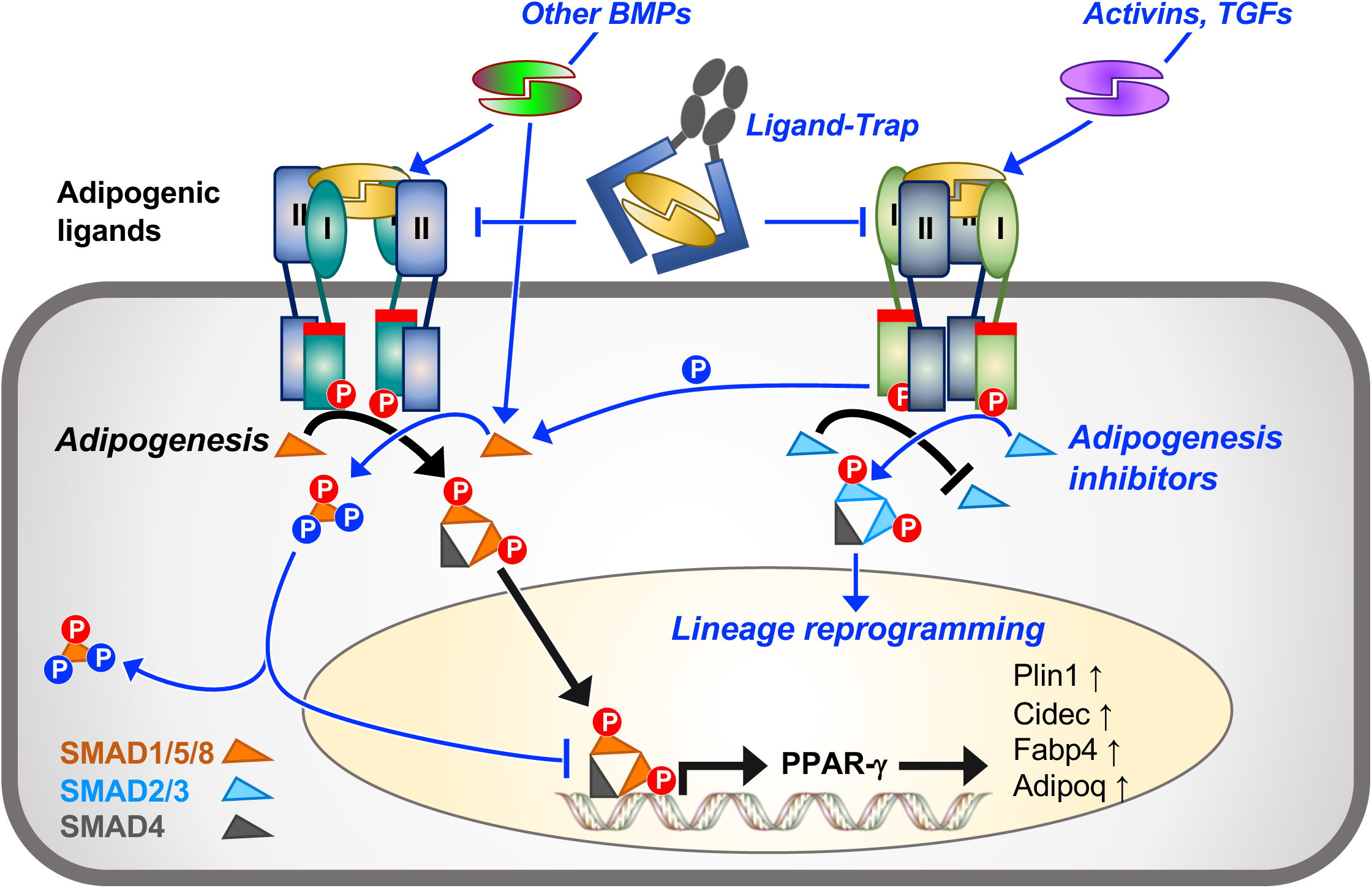
Model of SMAD regulation in adipogenesis and its inhibition. Mechanism of signaling during adipogenesis (represented by black arrows and lines). SMAD1/5/8 pathways (orange triangles) are activated as evidenced by C-terminal SMAD1/5/8 phosphorylation (red circled P) and SMAD1/5/8 dependent luciferase activity, leading to expression of adipogenic genes, including *Pparg*, *Plin1*, *Cidec*, *Fabp4* and *Adipoq*. By contrast, SMAD2/3 pathways (light blue triangles) are suppressed during adipogenesis. Signaling pathway reprograming following adipogenesis inhibitor treatment (represented by blue arrows and lines). Anti-adipogenic traps and ligands induce C-terminal SMAD2/3 phosphorylation (red circled P) and SMAD2/3 dependent luciferase activity. In parallel, SMAD1/5/8 is hyper-phosphorylation (blue circled P) by SMAD2/3 activating kinases or by SMAD2/3 regulated genes, leading to reduced SMAD1/5/8 nuclear translocation, SMAD1/5/8 transcriptional activities and adipogenic gene expression.

## DISCUSSION

Excessive caloric intake relative to energy expenditure stimulates adipose tissue (AT) expansion, a process that comprises adipocyte hypertrophy (increase in adipocyte size) and adipocyte hyperplasia (increase in adipocyte number) (Rutkowski et al., 2015). Adipocyte hyperplasia (a.k.a. adipogenesis) is triggered by signaling events that direct adipocyte precursors such as mesenchymal stem cells (MSCs) and adipose derived mesenchymal stromal cells (ADSC) to develop into mature adipocytes (Pittenger, Mackay et al., 1999, Rodeheffer, Birsoy et al., 2008). Adipogenic differentiation is of particular medical significance as ectopic fat deposition from newly formed adipocytes is strongly correlated with poor metabolic health (Fox, Massaro et al., 2007, Grundy, 2015, Jeffery, Church et al., 2015, Okamura, Hashimoto et al., 2018).

TGF-β family pathways feature prominently in adipogenesis (Lee, 2018, Tang & Lane, 2012, Zamani & Brown, 2011). As such, the family presents potential therapeutic targets that could help improve metabolic conditions by regulating formation of new adipocytes. However, TGF-β pathways are multi-functional, redundant and pleiotropic. Understanding how members of the family regulate adipocyte biology and how best to target these pathways in adipogenesis has, therefore, remained a challenge. We addressed this challenge using the 3T3-L1 adipogenesis model, as 3T3-L1 cells exhibit developmental and homeostatic properties of multiple adipocyte lineages (Morrison & McGee, 2015).

We first compared broadly the effect of various TGF-β family ligands on 3T3-L1 adipogenesis. Such a comprehensive approach provided robust, comparative data that helped us place individual activities within the broader context of TGF-β family signaling, allowing us to identify common themes in the mechanisms of adipogenic regulation by the family. Overall, we show that all SMAD2/3 pathway activating ligands can suppress adipogenesis and pre-adipocyte proliferation either by inhibiting precursor differentiation or, more likely, by reprograming precursors toward a non-adipogenic lineage. We also found that mature adipocytes became more hypertrophic in their presence, indicating that these ligands could potentially promote lipid storage or alter energy utilization by mature adipocytes. However, a reduced number of mature adipocytes exposed to high levels of nutrients could also explain the observed hypertrophy. By contrast, our results with SMAD1/5/8 pathway activating ligands were more subtle. As the SMAD1/5/8 pathway was activated in the basal state at the beginning of 3T3-L1 differentiation, we speculate that culture conditions provide key BMP growth factors or induce their expression to activate SMAD1/5/8 signaling and thus prime undifferentiated precursors for adipogenic commitment. Further addition of BMPs did not increase the level of 3T3-L1 cell commitment toward the adipogenic lineage as measured, *e.g.*, by an increased number of lipid droplets relative to the number of cells. However, we observed an overall increase in the number of lipid droplet-associated nuclei, indicating that BMP ligands may promote adipocyte hyperplasia by driving mitotic clonal expansion during differentiation. Notably, we did not find that BMP ligands promoted adipocyte hypertrophy as measured by an increase in lipid droplet size, lipid accumulation, or lipid droplet formation in treated cells. Taken together, these results clearly establish the distinct roles for each SMAD branch in adipogenesis and show that different ligands can adopt similar functions *in vitro*. These roles are largely based on the ligands’ ability to activate a particular SMAD branch. While a single ligand may prime all precursors for adipogenesis *in vivo,* the identity of the regulating ligand could also be specific to a particular adipogenic niche.

As one of our main goals was to identify inhibitors that prevent new fat cell formation, we adapted the 3T3-L1 adipogenesis model for *in vitro* screening of various ligand traps. These traps are engineered by fusing the ligand binding moieties of TGF-β family receptors, co-receptors or antagonists to an antibody Fc domain. They capture distinct groups of ligands in the extracellular space to block ligand-receptor binding and inhibit signaling. Significantly, we identified three traps that potently suppressed adipogenesis, as 3T3-L1 cells did not form lipid droplets or express adipogenic marker genes when treated with these traps. Intriguingly, TGFβRII-Fc was one of the most potent anti-adipogenic traps. This was unexpected as TGFβRII-Fc is well known as a specific inhibitor of the SMAD2/3 pathway activating ligands TGF-β1 and TGF-β3 (Aykul & Martinez-Hackert, 2016b), which also suppressed adipogenesis. Other adipogenesis inhibitors identified in this screen were Cripto-1-Fc and Cryptic-Fc, which inhibit Nodal and BMP-4 or Activin B, respectively (Aykul et al., 2017). Based on the binding specificities of the three anti-adipogenic traps, we could not single out one ligand that alone could account for their effect on adipocyte differentiation. In addition, we could not identify target ligands for these traps by pull-down and mass spectrometry in this assay, as their levels probably fall below the limit of detection. Our results, nevertheless, suggest that an interplay of multiple ligands or factors may be required to direct 3T3-L1 cells toward adipogenic fates. Notably, these results also suggest a complex mode of signal interpretation and processing by 3T3-L1 cells as both *bona fide* activators and inhibitors of SMAD2/3 signaling suppressed adipogenesis.

To elucidate the anti-adipogenic mechanism of the different traps and ligands, we investigated their effect on SMAD pathway activation and signal transduction. Using both anti p-SMAD Western blots and reporter gene expression assays, we showed that the SMAD1/5/8 pathway was activated in committed 3T3-L1 cells and further induced during their differentiation. To our surprise, all inhibitory traps and ligands, including the SMAD2/3 pathway activating ligand TGF-β1 and the TGF-β1 inhibitor trap TGFβRII-Fc, suppressed SMAD1/5/8 signaling as measured by reporter gene expression. As the SMAD1/5/8 signaling inhibitor LDN also prevented adipocyte formation, we propose that SMAD1/5/8 signaling is essential for adipogenesis. Strikingly, using anti p-SMAD Western blots we discovered that inhibitory traps or ligands did not suppress SMAD1/5/8 signaling by blocking C-terminal phosphorylation of SMAD1/5/8 as would be expected based on the canonical mechanism of TGF-β family signal transduction, and as we predictably detected in cells treated with the kinase inhibitor LDN. Instead, we consistently observed the near complete conversion of p-SMAD1/5/8 into a higher molecular weight p-SMAD1/5^Hi^ form that could be restored to its original electrophoretic mobility with alkaline phosphatase treatment. We, therefore, speculate that p-SMAD1/5^Hi^ represents hyper-phosphorylated SMAD1/5/8. In fact, all SMADs harbor a considerable number of kinase phosphorylation motifs that concentrate on their interdomain linker region (Figure S2), and phosphorylation of the SMAD linker has been demonstrated many times over and has been associated with various cellular outcomes, including both reduced and enhanced SMAD nuclear translocation and signaling (Alarcon et al., 2009, Blanchette, Rivard et al., 2001, Burch, Zheng et al., 2011, Gao, Alarcon et al., 2009, Kretzschmar, Doody et al., 1999). Indeed, we discovered that adipogenesis inhibitor treated cells, which predominantly exhibited hyper-phosphorylated p-SMAD1/5^Hi^, had greatly reduced SMAD1/5/8 signaling and did not differentiate into mature adipocytes. In addition, we mainly detected p-SMAD1/5^Hi^ in the cytoplasmic fraction, indicating that hyper-phosphorylation either prevented SMAD1/5/8 nuclear translocation, increased its export from the nucleus, or lead to its capture by ubiquitin ligases such as Smurf1, which could target SMAD1/5/8 for cytoplasmic retention or degradation (Kamato et al., 2013). However, as hyper-phosphorylation regulated pro-adipogenic SMAD1/5/8 signaling in 3T3-L1 cells, we propose that it presents an alternate, non-canonical mechanism of SMAD1/5/8 pathway regulation in adipogenesis that is distinct from the molecular step targeted by the small molecule kinase inhibitor LDN.

In contrast to SMAD1/5/8, we found that SMAD2/3 signaling was suppressed at all times during 3T3-L1 differentiation. Significantly, all tested traps and ligands that prevented adipogenesis activated SMAD2/3 signaling including, to our surprise, the *bona fide* SMAD2/3 signaling inhibitor TGFβRII-Fc. These findings demonstrate that SMAD2/3 activation is deleterious for adipogenic differentiation. Consistent with this interpretation, the SMAD2/3 activation inhibitor SB43 blunted the anti-adipogenic effect of inhibitory traps and ligands. Strikingly, SB43 treatment also suppressed formation of the p-SMAD1/5^Hi^ species and restored SMAD1/5/8 signaling, indicating that SMAD2/3 activation may be linked with SMAD1/5/8 inhibition by a regulatory feedback loop in adipocyte precursors. The proposed SMAD2/3 - SMAD1/5/8 feedback loop could be direct or indirect. *E.g.*, SB43 inhibited SMAD2/3 signaling, prevented formation of the p-SMAD1/5^Hi^ species and increased SMAD1/5/8 signaling, potentially implicating the SB43 target kinases ALK4, ALK5 or ALK7 directly in SMAD1/5/8 regulation and hyper-phosphorylation. Alternatively, SMAD2/3 pathway mediated expression or activation of intracellular kinases could provide an indirect mechanism of regulation. *E.g.*, hyper-phosphorylation of the SMAD linker region is suggested to serve as integration point for multiple signaling pathways to help modulate cellular responses to TGF-β family ligands (Massague, 2003), as expression or activation of intracellular kinases results in SMAD hyper-phosphorylation and inhibition of SMAD signaling (Hough et al., 2012, Kamato et al., 2013). Indeed, we observed increased phosphorylation (*i.e.*, activation) of the kinase ERK in cells treated with adipogenesis inhibitors. However, various ERK inhibitors failed to restore adipogenesis or prevent SMAD1/5/8 hyper-phosphorylation in cells treated with anti-adipogenic traps and ligands. Based on these findings, we speculate that SMAD1/5/8 hyper-phosphorylation could be mediated directly by an SB43 target kinase such as ALK4, ALK5, or ALK7 (Inman et al., 2002, Vogt, Traynor et al., 2011). Regardless of the specific mechanism of action that gives rise to p-SMAD1/5^Hi^, we identify a regulatory feedback loop that links SMAD2/3 activation with SMAD1/5/8 inhibition in adipogenesis. Notably, small molecule inhibition of SMAD1/5/8 did not alter SMAD2/3 activities, indicating that the feedback loop may only work in one direction.

In conclusion, we showed that SMAD1/5/8 pathways are activated while SMAD2/3 pathways are suppressed in differentiating 3T3-L1 cells. We identified several ligand traps that prevent 3T3-L1 adipogenesis via a conserved regulatory mechanism that involves differential activation of the SMAD2/3 and SMAD1/5/8 signaling branches by a negative feedback loop. Overall, these findings offer further insights into the complexity of TGF-β family action and regulation. Importantly, ligand traps with anti-adipogenic activity *in vitro* could help control hyperplastic AT expansion *in vivo*.

## METHODS

### Ligands

Human Activin A, Activin B, GDF-8, GDF-11, TGF-β1, BMP-2, BMP-4, BMP-6, BMP-9, and BMP-10 were obtained from R&D Systems, PROMOCELL or produced in-house. Activity was verified by Surface Plasmon Resonance and reporter gene assays.

### Fc-Fusion proteins

Synthetic genes of human ActRIIA, ActRIIB, ALK2, ALK3, ALK4, Cripto-1, and Cerberus, as well as mouse Cryptic were obtained from GeneArt. Human BMPRII and TGFβRII were PCR amplified from cDNA (Open Biosystems). Extracellular domains were fused to IgG1-Fc by PCR. Fc-fusion proteins were expressed using Chinese hamster ovary (CHO) cells, captured from condition medium using protein A affinity chromatography, eluted with 100 mM glycine, pH 3.0, and directly neutralized by adding 2% v/v 2 M Tris/HCl, pH 9.0. Purified proteins were either dialyzed directly into phosphate-buffered saline, pH 7.5, and stored at −80 °C or further purified by size exclusion chromatography in phosphate-buffered saline, pH 7.5, and stored at −80 °C. Purity was determined by SDS-PAGE and activity was verified by Surface Plasmon Resonance.

### Small molecule inhibitors

SB-431542 (SB43, 10 μM), LDN-193189 (LDN, 1 μM), PD98059 (PD98, 1 μM), PD0325901 (PD03, 1 μM), and the selective ERK inhibitor FR180204 (FR18, 10 μM) were purchased from Biovision. Samples were reconstituted in DMSO according to the manufactures’ instructions.

### 3T3-L1 differentiation assay

Murine 3T3-L1 pre-adipocytes were purchased from ZenBio. Cryopreserved 3T3-L1 cells were thawed and seeded at approximately 10,000 cells/cm^2^ in Preadipocyte Medium (PM: DMEM, high glucose HEPES pH 7.4, 10 % Bovine Calf Serum (BCS), and Penicillin + Streptomycin (PS)). Cells were maintained at 37°C in a humidified incubator with 5% CO_2_ until reaching 100% confluence (in about 4 days). During this time, media was replaced every other day. Two days after reaching confluence, Preadipocyte Medium (PM) was replaced with an appropriate volume of Differentiation Medium (DM: DMEM, high glucose, sodium pyruvate HEPES pH 7.4, 10 % Fetal Bovine Serum (FBS), 33 µM Biotin, 10 µg/ml Human insulin, 1 µM Dexamethasone, 0.5 mM 3-Isobutyl-1-methylxanthine (IBMX)) and incubated for 3 days. Differentiation Medium was then replaced with Adipocyte Maintenance Medium (MM: DMEM high glucose, sodium pyruvate HEPES pH 7.4, 10 % FBS, 33 µM Biotin, 10 µg/ml Human insulin,). Cells were maintained up to 10 days post differentiation with medium exchange every other day.

### Treatments

3T3-L1 cells were treated beginning at different stages of differentiation. Treatments were generally maintained until harvest. Briefly, confluent 3T3-L1cells were grown in PM, differentiated 3 days in DM and maintained up to 5 days in MM. 1 nM ligand (except TGF-β1, which is toxic, used at 0.1 nM) or 300 nM traps were added at day 0 (beginning of DM treatment), day 3 (end of DM treatment) or day 5 (after 2 days in MM). Cells were kept under treatment until the end of the experiment at day 8.

### Immunofluorescence

10,000 3T3-L1 cells/cm^2^ were plated in a 96-well plate in preadipocyte media. At day 0, cells were treated with differentiation medium containing test articles or small molecule inhibitors. Treatments were continued in the appropriate medium containing test articles as specified in the ‘3T3-L1 Differentiation assay’ section until day 10. At day 10, cells were washed twice with PBS and fixed with 10% formalin for 30 min at RT. Cells were then washed twice with PBS, followed by staining with 0.01% saponin, 1 µg/ml Nile Red and 1 µg/ml DAPI in PBS for 15 min at RT. After staining, cells were washed 3 times with PBS. Images were taken with Olympus Fluoview FC1000 confocal laser scanning microscope. In the figures, green represents Nile Red staining, purple represents DAPI staining. For quantitative Nile Red and DAPI fluorescence measurements, fluorescence was measured before and after staining according to published protocol (Aldridge, Kouroupis et al., 2013). Published data were obtained in duplicate. Assays were repeated multiple times.

### Image analysis

3T3-L1 cells were treated in quadruplicates in 96 well plate. Multiple images were taken from each well. Number of lipid droplets, number of nuclei, and mean lipid droplet intensity were calculated using ImageJ software from two biological replicates.

### qRT-PCR

Genomic DNA was isolated from 3T3-L1 cells and standard curves were created using a 1/3-fold dilution series with the highest concentration yielding qRT-PCR amplification at around cycle 20 for the majority of primer pairs. DNA and cDNA was quantified on a Lightcycler 480 (Roche) as described (Gjidoda et al., 2014). Primers were designed using the program PCRTiler (Gervais, Marques et al., 2010). A Tm-curve was performed as a quality control for each primer pair at the end of each qRT-PCR run to verify that only a single amplicon was produced by each primer pair and that no primer dimers were formed. Published data were obtained from two biological replicates.

### Immunoblots

∼19,000 3T3-L1 cells were plated in 24-well plates and grown to confluence in preadipocyte medium. Cells were then switched into differentiation and maintenance medium as required. Medium contained test articles, including ligands (0.1 - 1 nM), Fc-fusion traps constructs (−300 nM) and/or small molecule inhibitors (1 - 10 µM). After 3 to 10 days of cell growth at 37 °C, protein lysate was prepared by using ice-cold RIPA lysis buffer (150 mM NaCl, 1% Nonidet P-40, 0.1% SDS, 0.5% sodium deoxycholate, 50 mM Tris, pH 8.0, 1X “Protease Arrest” and 2X “Phosphatase Arrest” (G-Biosciences). Cell lysates were stored at −80 °C. Total protein concentration was determined by Bradford. For Western blot, equal amounts of protein (typically 10 μg) were separated under reducing conditions on 10 or 12 % TGX-polyacrylamide gels (Bio-Rad) and transferred to Hybond-P membrane (GE Healthcare). Membranes were blocked with Superblock (Thermofisher) and incubated with primary antibodies from Cell Signaling at a 1:1000 dilution (*e.g.* anti-phospho-SMAD2/3 (138D4), anti-phospho-SMAD1/5/8 (41D10)), or 1:5000 (*e.g.* anti-β-actin (8H10D10)), followed by incubation with horseradish peroxidase-conjugated secondary antibody at dilutions of 1:10000 (Actin) and 1:2000 (SMADs) dilution (7074). Western Bright ECL HRP substrate was used for detection (Advansta). Blots were visualized using autoradiography film. A list of antibodies can be found in the supplemental Table S3.

### Reporter gene expression

3T3-L1 cells in preadipocyte medium were seeded in each well of a 96-well plate and grown overnight. For transfection, solutions containing 0.2 µl/well Lipofectamine 3000, 0.2 µl/well P3000, 1 ng/well pGL4.74 plasmid (Luc2P/hRluc/TK, control luciferase reporter plasmid, Promega), and 100 ng/well of SMAD2/3 responsive reporter plasmid pGL4.48 (luc2P/SBE) or SMAD1/5/8 responsive reporter plasmid pGL3 (luc2P/BRE) in Opti-MEM were incubated at room temperature for 15 min. Transfection mixture was added to preadipocyte or differentiation medium containing test articles. Cells were then incubated for 16 h at 37 °C. Luciferase activity was detected using a homemade dual-glow luciferase assay (Baker & Boyce, 2014). Luminescence was determined using a FLUOstar Omega plate reader. Relative luciferase units were calculated by dividing firefly luciferase units with *Renilla* luciferase units (RLU). Data are expressed as mean of four independent measurements. Error bars correspond to S.E. of four independent measurements.

### Alkaline phosphatase treatment

440,000 3T3-L1 cells were plated in 100 mm dish and grown to confluence in preadipocyte medium. Cells were then switched into differentiation medium containing test articles at day 0. At day 3, cells were scraped in 50 mM Tris pH 8.0, 100 mM NaCl, 10 mM MgCl_2_, 1 mM DTT buffer and sonicated. Lysate concentration was determined by Bradford. 10 µg lysate was treated with 0.5 µl alkaline phosphatase (Sigma Aldrich, P0114) and incubated at 37 °C for 2 h. After incubation, samples were run on 10% TGX-polyacrylamide gels (Bio-Rad) and blotted with different antibodies.

### Cellular localization

Subcellular fractionation of 3T3-L1 cells was carried out according to published methods (https://bio-protocol.org/e754). Briefly, 440,000 3T3-L1 cells were plated in a 100 mm dish and grown to confluence in preadipocyte medium. Two days after plating, cells were switched into preadipocyte medium containing test articles. At day 0, cells were switched into differentiation medium containing test articles. Cells were harvested either at day 0 or day 3 in subcellular fractionation buffer (250mM sucrose, 20mM HEPES pH 7.4, 10mM KCl, 1.5 mm MgCl_2_, 1 mM EDTA,1 mM EGTA, 1 mM DTT, 1X “Protease Arrest” and 2X “Phosphatase Arrest” (G-biosciences). Cytoplasmic, nuclear and membrane fractions were separated as described (https://bio-protocol.org/e754). Equal total protein amounts of cytoplasmic, nuclear and membrane fractions were separated on 10% TGX gels (Bio-Rad) and blotted with different antibodies.

## ACKNOWLEDGEMENTS

We would like to thank Dr. Toril Holien for constructive comments on the manuscript. This work was funded in part by Michigan State University and by NIH grants R01 GM121499 to EMH.

## AUTHOR CONTRIBUTIONS

SA performed, SA and EMH designed and analyzed biochemistry, molecular biology and 3T3-L1 cell experiments; JM and MF designed, performed and analyzed qRT-PCR experiments; EMH wrote manuscript; SA, and EMH revised manuscript.

## COMPETING INTERESTS

All the work reported here was funded by federal grants or MSU funds. SA, EMH and/or MF have multiple patents (issued and pending) covering various ligand traps and their uses. EMH and SA hold options and/or shares of Talapo Therapeutics. EMH and MF are shareholders of Acceleron Pharma. SA is an employee and holds stock options of Regeneron Pharma. EMH and MF are founders, shareholders, and/or officers of Advertent Biotherapeutics. JM declares no competing interests.

## REFERENCES

1. Alarcon C, Zaromytidou AI, Xi Q, Gao S, Yu J, Fujisawa S, Barlas A, Miller AN, Manova-Todorova K, Macias MJ, Sapkota G, Pan D, Massague J (2009) Nuclear CDKs drive Smad transcriptional activation and turnover in BMP and TGF-beta pathways. Cell 139: 757–69

2. Aldridge A, Kouroupis D, Churchman S, English A, Ingham E, Jones E (2013) Assay validation for the assessment of adipogenesis of multipotential stromal cells--a direct comparison of four different methods. Cytotherapy 15: 89–101

3. Aubin J, Davy A, Soriano P (2004) In vivo convergence of BMP and MAPK signaling pathways: impact of differential Smad1 phosphorylation on development and homeostasis. Genes Dev 18: 1482–94

4. Aykul S, Martinez-Hackert E (2016a) New Ligand Binding Function of Human Cerberus and Role of Proteolytic Processing in Regulating Ligand-Receptor Interactions and Antagonist Activity. J Mol Biol 428: 590–602

5. Aykul S, Martinez-Hackert E (2016b) Transforming Growth Factor-beta Family Ligands Can Function as Antagonists by Competing for Type II Receptor Binding. J Biol Chem 291: 10792–804

6. Aykul S, Parenti A, Chu KY, Reske J, Floer M, Ralston A, Martinez-Hackert E (2017) Biochemical and Cellular Analysis Reveals Ligand Binding Specificities, a Molecular Basis for Ligand Recognition, and Membrane Association-dependent Activities of Cripto-1 and Cryptic. J Biol Chem 292: 4138–4151

7. Baker JM, Boyce FM (2014) High-throughput functional screening using a homemade dual-glow luciferase assay. Journal of visualized experiments: JoVE Baud’huin M, Solban N, Cornwall-Brady M, Sako D, Kawamoto Y, Liharska K, Lath D, Bouxsein ML, Underwood KW, Ucran J, Kumar R, Pobre E, Grinberg A, Seehra J, Canalis E, Pearsall RS, Croucher PI (2012) A soluble bone morphogenetic protein type IA receptor increases bone mass and bone strength. Proc Natl Acad Sci U S A 109: 12207–12

8. Blanchette F, Rivard N, Rudd P, Grondin F, Attisano L, Dubois CM (2001) Cross-talk between the p42/p44 MAP kinase and Smad pathways in transforming growth factor beta 1-induced furin gene transactivation. J Biol Chem 276: 33986–94

9. Bost F, Aouadi M, Caron L, Binetruy B (2005) The role of MAPKs in adipocyte differentiation and obesity. Biochimie 87: 51–6

10. Bowers RR, Lane MD (2007) A role for bone morphogenetic protein-4 in adipocyte development. Cell Cycle 6: 385–9

11. Burch ML, Zheng W, Little PJ (2011) Smad linker region phosphorylation in the regulation of extracellular matrix synthesis. Cellular and molecular life sciences: CMLS 68: 97–107

12. Choy L, Skillington J, Derynck R (2000) Roles of autocrine TGF-beta receptor and Smad signaling in adipocyte differentiation. J Cell Biol 149: 667–82

13. Cuny GD, Yu PB, Laha JK, Xing X, Liu JF, Lai CS, Deng DY, Sachidanandan C, Bloch KD, Peterson RT (2008) Structure-activity relationship study of bone morphogenetic protein (BMP) signaling inhibitors. Bioorg Med Chem Lett 18: 4388–92

14. Daly AC, Randall RA, Hill CS (2008) Transforming growth factor beta-induced Smad1/5 phosphorylation in epithelial cells is mediated by novel receptor complexes and is essential for anchorage-independent growth. Mol Cell Biol 28: 6889–902

15. David L, Mallet C, Mazerbourg S, Feige JJ, Bailly S (2007) Identification of BMP9 and BMP10 as functional activators of the orphan activin receptor-like kinase 1 (ALK1) in endothelial cells. Blood 109: 1953–61

16. de Caestecker M (2004) The transforming growth factor-beta superfamily of receptors. Cytokine Growth Factor Rev 15: 1–11

17. Derynck R, Zhang YE (2003) Smad-dependent and Smad-independent pathways in TGF-beta family signalling. Nature 425: 577–84

18. Eckel RH, Grundy SM, Zimmet PZ (2005) The metabolic syndrome. Lancet 365: 1415–28

19. Fox CS, Massaro JM, Hoffmann U, Pou KM, Maurovich-Horvat P, Liu CY, Vasan RS, Murabito JM, Meigs JB, Cupples LA, D’Agostino RB, Sr., O’Donnell CJ (2007) Abdominal visceral and subcutaneous adipose tissue compartments: association with metabolic risk factors in the Framingham Heart Study. Circulation 116: 39–48

20. Fuentealba LC, Eivers E, Ikeda A, Hurtado C, Kuroda H, Pera EM, De Robertis EM (2007) Integrating patterning signals: Wnt/GSK3 regulates the duration of the BMP/Smad1 signal. Cell 131: 980–93

21. Gao S, Alarcon C, Sapkota G, Rahman S, Chen PY, Goerner N, Macias MJ, Erdjument-Bromage H, Tempst P, Massague J (2009) Ubiquitin ligase Nedd4L targets activated Smad2/3 to limit TGF-beta signaling. Mol Cell 36: 457–68

22. Gervais AL, Marques M, Gaudreau L (2010) PCRTiler: automated design of tiled and specific PCR primer pairs. Nucleic acids research 38: W308–12

23. Gjidoda A, Tagore M, McAndrew MJ, Woods A, Floer M (2014) Nucleosomes are stably evicted from enhancers but not promoters upon induction of certain pro-inflammatory genes in mouse macrophages. PLoS One 9: e93971

24. Grundy SM (2015) Adipose tissue and metabolic syndrome: too much, too little or neither. European journal of clinical investigation 45: 1209–17

25. Gulden E, Mollerus S, Bruggemann J, Burkart V, Habich C (2008) Heat shock protein 60 induces inflammatory mediators in mouse adipocytes. FEBS Lett 582: 2731–6

26. Gustafson B, Hammarstedt A, Hedjazifar S, Hoffmann JM, Svensson PA, Grimsby J, Rondinone C, Smith U (2015) BMP4 and BMP Antagonists Regulate Human White and Beige Adipogenesis. Diabetes 64: 1670–81

27. Hamza MS, Pott S, Vega VB, Thomsen JS, Kandhadayar GS, Ng PW, Chiu KP, Pettersson S, Wei CL, Ruan Y, Liu ET (2009) De-novo identification of PPARgamma/RXR binding sites and direct targets during adipogenesis. PLoS One 4: e4907

28. Hayashida T, Decaestecker M, Schnaper HW (2003) Cross-talk between ERK MAP kinase and Smad signaling pathways enhances TGF-beta-dependent responses in human mesangial cells. Faseb J 17: 1576–8

29. Herrera B, van Dinther M, Ten Dijke P, Inman GJ (2009) Autocrine bone morphogenetic protein-9 signals through activin receptor-like kinase-2/Smad1/Smad4 to promote ovarian cancer cell proliferation. Cancer Res 69: 9254–62

30. Hirai S, Yamanaka M, Kawachi H, Matsui T, Yano H (2005) Activin A inhibits differentiation of 3T3-L1 preadipocyte. Mol Cell Endocrinol 232: 21–6

31. Hoggard N, Cruickshank M, Moar KM, Barrett P, Bashir S, Miller JD (2009) Inhibin betaB expression in murine adipose tissue and its regulation by leptin, insulin and dexamethasone. Journal of molecular endocrinology 43: 171–7

32. Hough C, Radu M, Dore JJ (2012) Tgf-beta induced Erk phosphorylation of smad linker region regulates smad signaling. PLoS One 7: e42513

33. Huang H, Song TJ, Li X, Hu L, He Q, Liu M, Lane MD, Tang QQ (2009) BMP signaling pathway is required for commitment of C3H10T1/2 pluripotent stem cells to the adipocyte lineage. Proc Natl Acad Sci U S A 106: 12670–5

34. Ignotz RA, Massague J (1985) Type beta transforming growth factor controls the adipogenic differentiation of 3T3 fibroblasts. Proc Natl Acad Sci U S A 82: 8530–4

35. Inman GJ, Nicolas FJ, Callahan JF, Harling JD, Gaster LM, Reith AD, Laping NJ, Hill CS (2002) SB-431542 is a potent and specific inhibitor of transforming growth factor-beta superfamily type I activin receptor-like kinase (ALK) receptors ALK4, ALK5, and ALK7. Mol Pharmacol 62: 65-74

36. Jeffery E, Church CD, Holtrup B, Colman L, Rodeheffer MS (2015) Rapid depot-specific activation of adipocyte precursor cells at the onset of obesity. Nat Cell Biol 17: 376–85

37. Kamato D, Burch ML, Piva TJ, Rezaei HB, Rostam MA, Xu S, Zheng W, Little PJ, Osman N (2013) Transforming growth factor-beta signalling: role and consequences of Smad linker region phosphorylation. Cell Signal 25: 2017–24

38. Kaplan NM (1989) The deadly quartet. Upper-body obesity, glucose intolerance, hypertriglyceridemia, and hypertension. Archives of internal medicine 149: 1514–20

39. Kim HS, Liang L, Dean RG, Hausman DB, Hartzell DL, Baile CA (2001) Inhibition of preadipocyte differentiation by myostatin treatment in 3T3-L1 cultures. Biochem Biophys Res Commun 281: 902–6

40. Koster J, Volckmann R, Zwijnenburg D, Molenaar P, Versteeg R (2019) R2: Genomics analysis and visualization platform. Cancer Res 79

41. Kretzschmar M, Doody J, Massague J (1997) Opposing BMP and EGF signalling pathways converge on the TGF-beta family mediator Smad1. Nature 389: 618–22

42. Kretzschmar M, Doody J, Timokhina I, Massague J (1999) A mechanism of repression of TGFbeta/ Smad signaling by oncogenic Ras. Genes Dev 13: 804–16

43. Kuroda H, Fuentealba L, Ikeda A, Reversade B, De Robertis EM (2005) Default neural induction: neuralization of dissociated Xenopus cells is mediated by Ras/MAPK activation. Genes Dev 19: 1022–7

44. Lattin JE, Schroder K, Su AI, Walker JR, Zhang J, Wiltshire T, Saijo K, Glass CK, Hume DA, Kellie S, Sweet MJ (2008) Expression analysis of G Protein-Coupled Receptors in mouse macrophages. Immunome Res 4: 5

45. Lee MJ (2018) Transforming growth factor beta superfamily regulation of adipose tissue biology in obesity. Biochim Biophys Acta 1864: 1160–1171

46. Lee MJ, Pickering RT, Shibad V, Wu Y, Karastergiou K, Jager M, Layne MD, Fried SK (2019) Impaired Glucocorticoid Suppression of TGFbeta Signaling in Human Omental Adipose Tissues Limits Adipogenesis and May Promote Fibrosis. Diabetes 68: 587–597

47. Lewis KA, Gray PC, Blount AL, MacConell LA, Wiater E, Bilezikjian LM, Vale W (2000) Betaglycan binds inhibin and can mediate functional antagonism of activin signalling. Nature 404: 411–4

48. Luo H, Guo Y, Liu Y, Wang Y, Zheng R, Ban Y, Peng L, Yuan Q, Liu W (2019) Growth differentiation factor 11 inhibits adipogenic differentiation by activating TGF-beta/Smad signalling pathway. Cell Prolif 52: e12631

49. Massague J (2003) Integration of Smad and MAPK pathways: a link and a linker revisited. Genes Dev 17: 2993–7

50. Matsuura I, Wang G, He D, Liu F (2005) Identification and characterization of ERK MAP kinase phosphorylation sites in Smad3. Biochemistry 44: 12546–53

51. Modica S, Wolfrum C (2017) The dual role of BMP4 in adipogenesis and metabolism. Adipocyte 6: 141–146

52. Morrison S, McGee SL (2015) 3T3-L1 adipocytes display phenotypic characteristics of multiple adipocyte lineages. Adipocyte 4: 295–302

53. Moustakas A, Souchelnytskyi S, Heldin CH (2001) Smad regulation in TGF-beta signal transduction. J Cell Sci 114: 4359–69

54. Nakao A, Imamura T, Souchelnytskyi S, Kawabata M, Ishisaki A, Oeda E, Tamaki K, Hanai J, Heldin CH, Miyazono K, ten Dijke P (1997) TGF-beta receptor-mediated signalling through Smad2, Smad3 and Smad4. Embo J 16: 5353-62

55. Okamura T, Hashimoto Y, Hamaguchi M, Obora A, Kojima T, Fukui M (2018) Ectopic fat obesity presents the greatest risk for incident type 2 diabetes: a population-based longitudinal study. International journal of obesity

56. Olsen OE, Sankar M, Elsaadi S, Hella H, Buene G, Darvekar SR, Misund K, Katagiri T, Knaus P, Holien T (2018) BMPR2 inhibits activin and BMP signaling via wild-type ALK2. J Cell Sci 131

57. Olsen OE, Wader KF, Misund K, Vatsveen TK, Ro TB, Mylin AK, Turesson I, Stordal BF, Moen SH, Standal T, Waage A, Sundan A, Holien T (2014) Bone morphogenetic protein-9 suppresses growth of myeloma cells by signaling through ALK2 but is inhibited by endoglin. Blood Cancer J 4: e196

58. Pera EM, Ikeda A, Eivers E, De Robertis EM (2003) Integration of IGF, FGF, and anti-BMP signals via Smad1 phosphorylation in neural induction. Genes Dev 17: 3023-8

59. Pittenger MF, Mackay AM, Beck SC, Jaiswal RK, Douglas R, Mosca JD, Moorman MA, Simonetti DW, Craig S, Marshak DR (1999) Multilineage potential of adult human mesenchymal stem cells. Science 284: 143–7

60. Rebbapragada A, Benchabane H, Wrana JL, Celeste AJ, Attisano L (2003) Myostatin signals through a transforming growth factor beta-like signaling pathway to block adipogenesis. Mol Cell Biol 23: 7230–42

61. Rodeheffer MS, Birsoy K, Friedman JM (2008) Identification of white adipocyte progenitor cells in vivo. Cell 135: 240–9

62. Rosen ED (2005) The transcriptional basis of adipocyte development. Prostaglandins Leukot Essent Fatty Acids 73: 31–4

63. Rutkowski JM, Stern JH, Scherer PE (2015) The cell biology of fat expansion. J Cell Biol 208: 501–12

64. Sako D, Grinberg AV, Liu J, Davies MV, Castonguay R, Maniatis S, Andreucci AJ, Pobre EG, Tomkinson KN, Monnell TE, Ucran JA, Martinez-Hackert E, Pearsall RS, Underwood KW, Seehra J, Kumar R (2010) Characterization of the ligand binding functionality of the extracellular domain of activin receptor type IIb. J Biol Chem 285: 21037–48

65. Sapkota G, Alarcon C, Spagnoli FM, Brivanlou AH, Massague J (2007) Balancing BMP signaling through integrated inputs into the Smad1 linker. Mol Cell 25: 441–54

66. Scharpfenecker M, van Dinther M, Liu Z, van Bezooijen RL, Zhao Q, Pukac L, Lowik CW, ten Dijke P (2007) BMP-9 signals via ALK1 and inhibits bFGF-induced endothelial cell proliferation and VEGF-stimulated angiogenesis. J Cell Sci 120: 964–72

67. Schreiber I, Dorpholz G, Ott CE, Kragesteen B, Schanze N, Lee CT, Kohrle J, Mundlos S, Ruschke K, Knaus P (2017) BMPs as new insulin sensitizers: enhanced glucose uptake in mature 3T3-L1 adipocytes via PPARgamma and GLUT4 upregulation. Scientific reports 7: 17192

68. Shi Y, Massague J (2003) Mechanisms of TGF-beta signaling from cell membrane to the nucleus. Cell 113: 685–700

69. Sparks RL, Allen BJ, Strauss EE (1992) TGF-beta blocks early but not late differentiation-specific gene expression and morphologic differentiation of 3T3 T proadipocytes. J Cell Physiol 150: 568–77

70. Stephens JM, Morrison RF, Wu Z, Farmer SR (1999) PPARgamma ligand-dependent induction of STAT1, STAT5A, and STAT5B during adipogenesis. Biochem Biophys Res Commun 262: 216-22

71. Suenaga M, Kurosawa N, Asano H, Kanamori Y, Umemoto T, Yoshida H, Murakami M, Kawachi H, Matsui T, Funaba M (2013) Bmp4 expressed in preadipocytes is required for the onset of adipocyte differentiation. Cytokine 64: 138–45

72. Sun K, Kusminski CM, Scherer PE (2011) Adipose tissue remodeling and obesity. J Clin Invest 121: 2094–101

73. Sun K, Scherer PE (2010) Adipose Tissue Dysfunction: A Multistep Process. Res Perspect End Int: 67–75

74. Tang QQ, Lane MD (2012) Adipogenesis: from stem cell to adipocyte. Annual review of biochemistry 81: 715–36

75. Tang QQ, Otto TC, Lane MD (2004) Commitment of C3H10T1/2 pluripotent stem cells to the adipocyte lineage. Proc Natl Acad Sci U S A 101: 9607–11

76. Townson SA, Martinez-Hackert E, Greppi C, Lowden P, Sako D, Liu J, Ucran JA, Liharska K, Underwood KW, Seehra J, Kumar R, Grinberg AV (2012) Specificity and Structure of a High Affinity Activin Receptor-like Kinase 1 (ALK1) Signaling Complex. J Biol Chem 287: 27313–25

77. Tseng YH, Kokkotou E, Schulz TJ, Huang TL, Winnay JN, Taniguchi CM, Tran TT, Suzuki R, Espinoza DO, Yamamoto Y, Ahrens MJ, Dudley AT, Norris AW, Kulkarni RN, Kahn CR (2008) New role of bone morphogenetic protein 7 in brown adipogenesis and energy expenditure. Nature 454: 1000–4

78. Vogt J, Traynor R, Sapkota GP (2011) The specificities of small molecule inhibitors of the TGFss and BMP pathways. Cell Signal 23: 1831–42

79. von Bubnoff A, Peiffer DA, Blitz IL, Hayata T, Ogata S, Zeng Q, Trunnell M, Cho KW (2005) Phylogenetic footprinting and genome scanning identify vertebrate BMP response elements and new target genes. Dev Biol 281: 210–26

80. Walker RG, Czepnik M, Goebel EJ, McCoy JC, Vujic A, Cho M, Oh J, Aykul S, Walton KL, Schang G, Bernard DJ, Hinck AP, Harrison CA, Martinez-Hackert E, Wagers AJ, Lee RT, Thompson TB (2017) Structural basis for potency differences between GDF8 and GDF11. BMC biology 15: 19

81. Wang QA, Tao C, Gupta RK, Scherer PE (2013) Tracking adipogenesis during white adipose tissue development, expansion and regeneration. Nature medicine 19: 1338–44

82. Zamani N, Brown CW (2011) Emerging roles for the transforming growth factor-{beta} superfamily in regulating adiposity and energy expenditure. Endocrine reviews 32: 387–403

83. Zebisch K, Voigt V, Wabitsch M, Brandsch M (2012) Protocol for effective differentiation of 3T3-L1 cells to adipocytes. Analytical biochemistry 425: 88–90

